# An 11-point time course midgut transcriptome across 72 h after blood feeding provides detailed temporal resolution of transcript expression in the arbovirus vector, *Aedes aegypti*

**DOI:** 10.1101/2023.03.03.531062

**Authors:** Hitoshi Tsujimoto, Zach N. Adelman

**Author notes:** Corresponding author Zach N Adelman 329A Minnie Belle Heep Center Dept of Entomology 370 Olsen Blvd Texas A&M University College Station, TX 77843.

## Abstract

As the major vector for dengue, Zika, yellow fever, and chikungunya viruses, the mosquito *Aedes aegypti* is one of the most important insects in public health. These viruses are transmitted by bloodfeeding, which is also necessary for the reproduction of the mosquito. Thus, the midgut plays an essential role in mosquito physiology as the center for bloodmeal digestion and as an organ that serves as the first line of defense against viruses. Despite its importance, transcriptomic dynamics with fine temporal resolution across the entire digestion cycle have not yet been reported. To fill this gap, we conducted a transcriptomic analysis of *Ae. aegypti* female midgut across a 72-h bloodmeal digestion cycle for 11 time points with a particular focus on the first 24 h. PCA analysis confirmed that 72 h is indeed a complete digestion cycle. Cluster and GO enrichment analysis showed the orchestrated modulation of thousands of genes to accomplish the midgut’s role as the center for digestion as well as nutrient transport with a clear progression with sequential emphasis on transcription, translation, energy production, nutrient metabolism, transport, and finally autophagy by 24-36hr. We further determined that many serine proteases are robustly expressed as if to prepare for unexpected physiological challenges. This study provides a powerful resource for the analysis of genomic features that coordinate the rapid and complex transcriptional program induced by mosquito bloodfeeding.

## Introduction

Arthropod-borne (arbo-) viruses impose a substantial burden on public health (WHO 2009; WHO 2016; WHO 2018; WHO 2019). As the major vector of dengue, Zika, yellow fever and chikungunya viruses, the mosquito *Aedes aegypti* plays a leading role in arbovirus transmission. Transmission of the viruses occur during bloodfeeding, which is required for reproduction of the mosquito. An *Ae. aegypti* female may take ∼2.5 mg of blood, which is ∼219% of its body weight (Stobbart 1977) and produces 60-150 eggs per batch. Laboratory strains of *Ae. aegypti* perform this feat (from blood ingestion to maturation of ovaries) in ∼72 h. In the meantime, blood-acquired pathogens can invade midgut epithelial cells despite the presence of physical and biochemical barriers, replicate, and disseminate throughout the body of the mosquito, including the salivary glands from which the pathogens will be injected into a new vertebrate host at the next bloodfeeding. Thus, the female midgut is one of the most important organs in a mosquito as the center for bloodmeal digestion, which is directly related to reproduction and the first line of defense against blood-acquired pathogens.

Therefore, a detailed understanding of midgut physiology and the underlying genetic basis for major aspects of digestion and reproduction should provide extremely valuable knowledge to help identify novel avenues of preventing pathogen transmission or of killing bloodfeeding mosquitoes. Among many methods, transcriptomic approaches provide a global view of expressed genes in a tissue or organism, which has been useful to determine previously unrecognized genes to exploit for novel control strategies. Previous studies of the *Ae. aegypti* midgut transcriptome demonstrated that the midgut undergoes very complex control of gene expression in response not only to a bloodmeal, but also to pathogens, which may be present in the bloodmeal (Bonizzoni et al. 2012b; Bonizzoni et al. 2012a). Recent studies have added spatial resolution to the female midgut transcriptome. Hixson *et al*. revealed that the subsections of midgut represent quite different transcriptome profiles (Hixson et al. 2022); and another study utilizing single-nuclei RNAseq uncovered that the midgut is comprised of at least 20 different cell types (Cui and Franz 2020). Moreover, Raquin *et al*. analyzed individual midgut RNAseq to determine host factors important for dengue virus (DENV) replication (Raquin et al. 2017), which added resolution at individual level.

A female *Ae. aegypti* undergoes dynamic physiological change in response to a bloodmeal. For instance, about 24% of the bloodmeal weight will be excreted as urine within 30 min after ingestion (Stobbart 1977). Formation of the peritrophic matrix (PM), an extracellular layer that forms around the bloodmeal to protect midgut epithelium and consists of chitin and associated proteins also appears to occur strikingly quick: a structure stained by azan (a stain for fibrous connective materials) began to form between the midgut epithelium and bloodmeal at 50 min post bloodmeal (PBM) (Freyvogel and Staeubli 1965), while expression of two major peritrophic matrix proteins (peritrophins) have been induced within 5 h after bloodmeal uptake (Shao et al. 2005; Devenport et al. 2006). Although the midgut is expected to play a major role in such physiological changes, prior studies to date have covered only a few time points, which were mostly relevant to virus infection cycle (extrinsic incubation period of dengue virus: 6.5 to 15 days at 30 and 25 °C, respectively (Chan and Johansson 2012)): 5 h in (Bonizzoni et al. 2011); 1, 4 and 14 days in (Bonizzoni et al. 2012a); 1 and 4 days in (Raquin et al. 2017); 4-6, 24, 48 h (of the “gut” including crop, midgut and hindgut) in (Hixson et al. 2022). One study analyzed a 72-h time course over 5 time points (0, 3, 12, 24, 72 hPBM) using the hybridization-based microarray technology (Sanders et al. 2003). However, the number of genes it could analyze (pre-determined 1778 genes) were far from modern sequencing-based technologies in quantity and quality, which makes a fair comparison unreasonable. Thus, the precise timing of changes in the midgut transcriptome over the course of bloodmeal digestion, especially within 24 h remains largely unexplored.

To fill the gap and to uncover the breadth of dynamic change in the transcriptome throughout the digestion cycle, we conducted a transcriptomic study for 11 time points across 72 h upon bloodfeeding, which should serve as a platform for planning experiments on *Ae. aegypti* bloodmeal digestion and reproduction physiology at genetic and genomic viewpoints. Such studies will stimulate further advancement of *Ae. aegypti* research.

## Results

### Morphology of midgut during the time-course

We captured images of the midgut with associated organs (Malpighian tubules, hindgut and ovaries) (Fig 1, Supplemental Fig S1). These images depict a midgut digestion cycle (pre-bloodmeal and from uptake of a bloodmeal to complete clearance of the midgut) that takes 48-72 h, supporting the idea that our time course should provide with a fairly complete view of the change in transcript expression throughout the digestion cycle.

**Fig 1:**
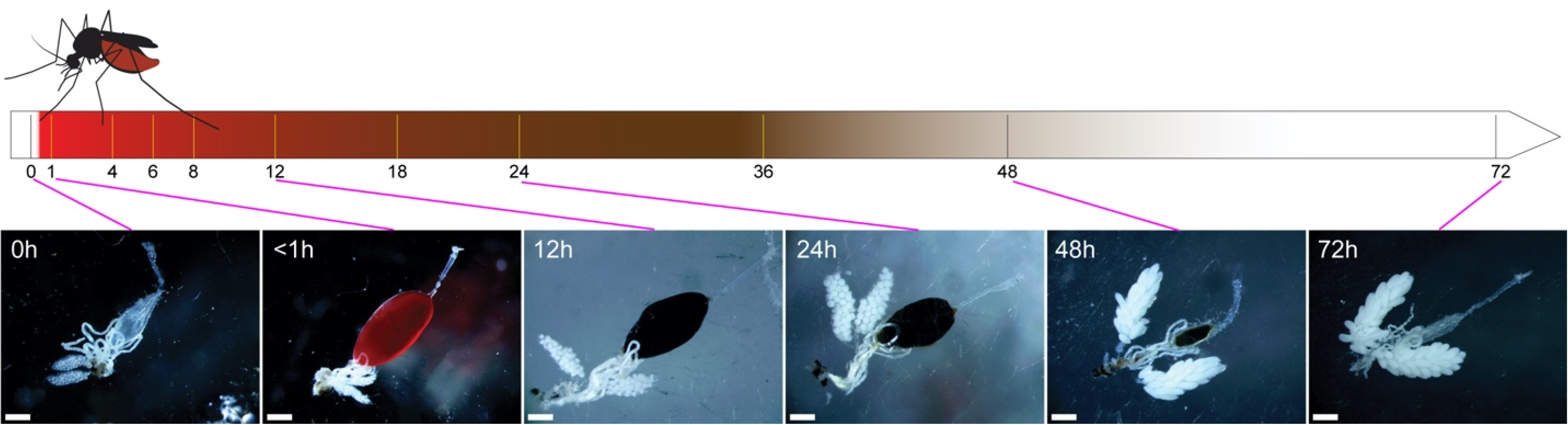
Timeline for the midgut transcriptome sample collection time points with images of the gut system and ovaries for selected time points. Images of all the time points are shown in Supplemental Fig S1. Scale bar (white bar in each panel): 500 _μ_m.

### Sequencing experiments

We utilized a laboratory strain of *Ae. aegypti* “Liverpool”, which has a long history of usage in research and is a reference strain for the genome sequencing (Kuno 2010; Matthews et al. 2018). For the experiments for transcriptome sequencing, we used citrated sheep blood instead of defibrinated sheep blood, as the latter is not whole blood and we considered that the response of the mosquitoes may be more natural with whole blood (such as citrated blood). We performed dissection of the midguts at the 11 time points for light-cycle matched female mosquitoes (30 midguts per time point for a replicate). To verify the consistency of each experiment, we replicated the experiment independently four times. RNA extraction, library prep, and Illumina sequencing are detailed in the Materials and Methods.

### RNAseq output and mapping

From our 4 independent replicates for the 11 time points of pre- and post-bloodmeal, Illumina NovaSeq 6000 paired end sequencing generated 1,160,607,602 fragment pairs (= 2,321,215,204 reads). We obtained a mean of 26,377,445.5 fragments per sample (ranging 19,503,828 to 34,008,005 fragments) and 105,509,782 fragments per time point. We mapped these fragments using HISAT2 (ver. 2.1.0) (Pertea et al. 2016) onto the AaegL5 AGWG genome assembly (obtained from VectorBase: https://vectorbase.org/vectorbase/app) (Giraldo-Calderón et al. 2015; Matthews et al. 2018) with 78.48% average mapping rate (Supplemental Fig S2).

Through quantification of expression by StringTie (ver. 2.0) (Pertea et al. 2016), we detected 16,535 genes with non-zero TPM values in the midgut, of which 13,162 were protein coding genes (89.4% of total in the genome), 3,134 non-coding RNA genes (66.6%), and 260 pseudogenes (68.0%) from the current gene annotation (AaegL5.3).

### Principal component analysis (PCA)

As a process of validation for consistency of the replicates and changes in gene expression between time points, we performed principal component analysis (PCA) by DESeq2 (ver. 1.34.0) (Love et al. 2014) (Fig 2). PCA showed that the four replicates at all the timepoints were reasonably clustered, suggesting consistency between experiments. The expression changes across the time points showed a circular trace. Surprisingly, 392 genes were differentially expressed even within <1 hPBM compared to the pre-bloodfed control (by DESeq2) (Supplemental Table S1). We term it as <1 hPBM for simplicity, but it was in fact around 15-20 min after bloodfeeding (see Materials and Methods), which suggests that the response of the female midgut to a bloodmeal is strikingly rapid. The difference between <1 and 4 hPBM appeared to show the largest difference between time points (5115 genes, the highest number between neighboring time points, Table S2), which also suggests a rapid change in transcription in the first 4 h. The timepoints that showed the greatest difference from the pre-bloodmeal state (0 hPBM) were 18-24 hPBM, and the expression profile turned to near 0 hPBM at 72 hPBM (Fig 2).

**Fig 2:**
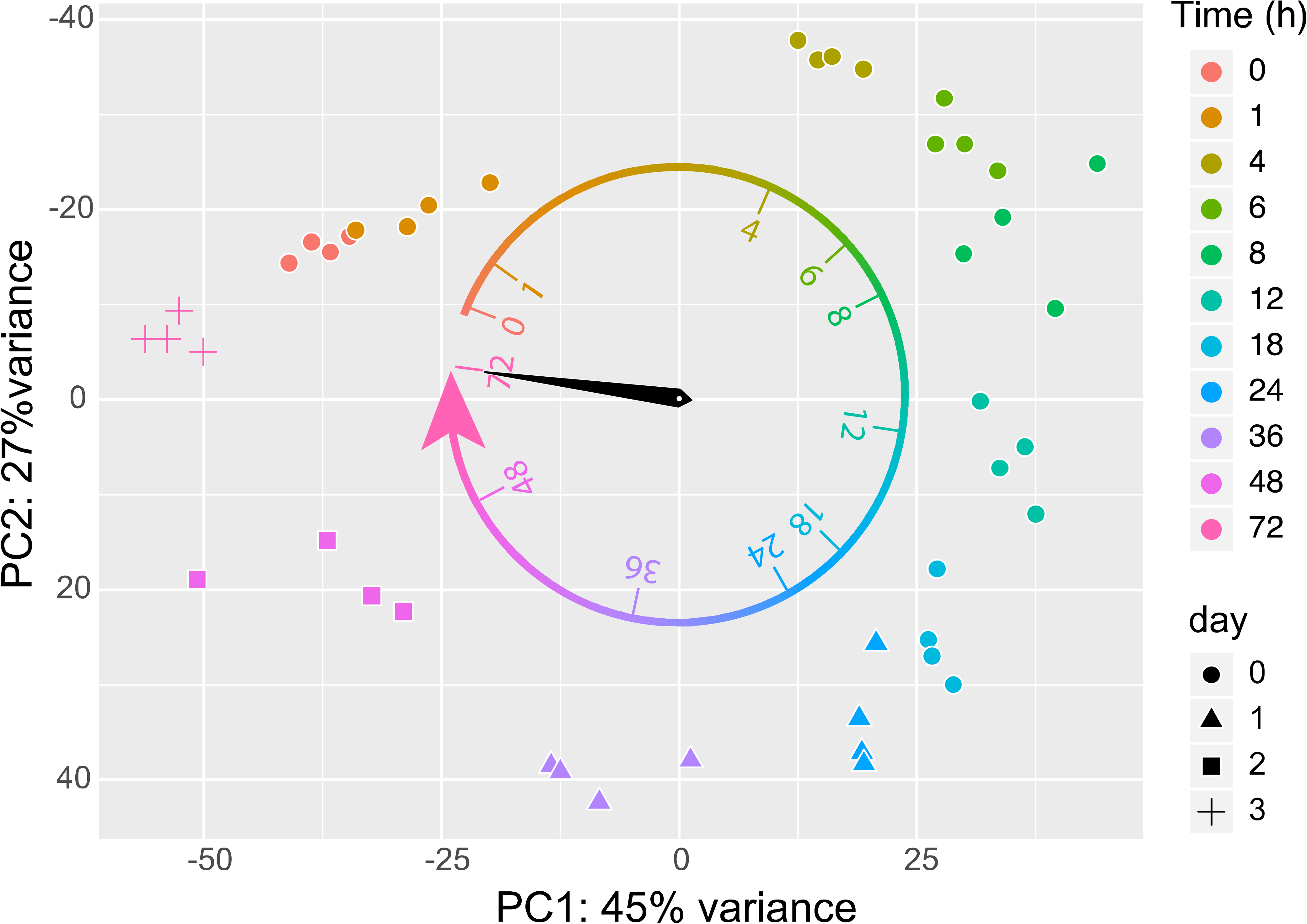
Principal component analysis (PCA) of the libraries. Time points (hPBM) are shown by the color scheme and the point shapes. The circular arrow with a clock hand indicates a 72-h cycle.

### Time-course analysis and Mfuzz soft clustering

Through edgeR (ver. 3.36.0) (Chen et al. 2016) time-course analysis, 8594 genes showed significant change at FDR < 0.05. These genes were clustered by Mfuzz (2.54.0) (Futschik and Carlisle 2005) into 20 clusters, which were manually consolidated into 13 groups by similar expression patterns, which were named by the expression peaks and bottoms: 1) 4h peak, 2) 6h peak, 3) 8h peak, 4) 12h peak, 5) 18-24h peak, 6) 24-36h peak, 7) 36h peak, 8) 48h peak, 9) up (consistently elevated expression relative to pre-bloodmeal state), 10) 6h bottom, 11) 18h bottom, 12) 48h bottom, and 13) down (Supplemental Fig S3).

Gene ontology (GO) enrichment analyses using analysis tool in the VectorBase website (Giraldo-Calderón et al. 2015), with a primary focus on “Biological Process” (BP) terms, indicated a far greater numbers of terms enriched in the first three peak groups (Fig 3 and Supplemental Table S2).

**Fig3:**
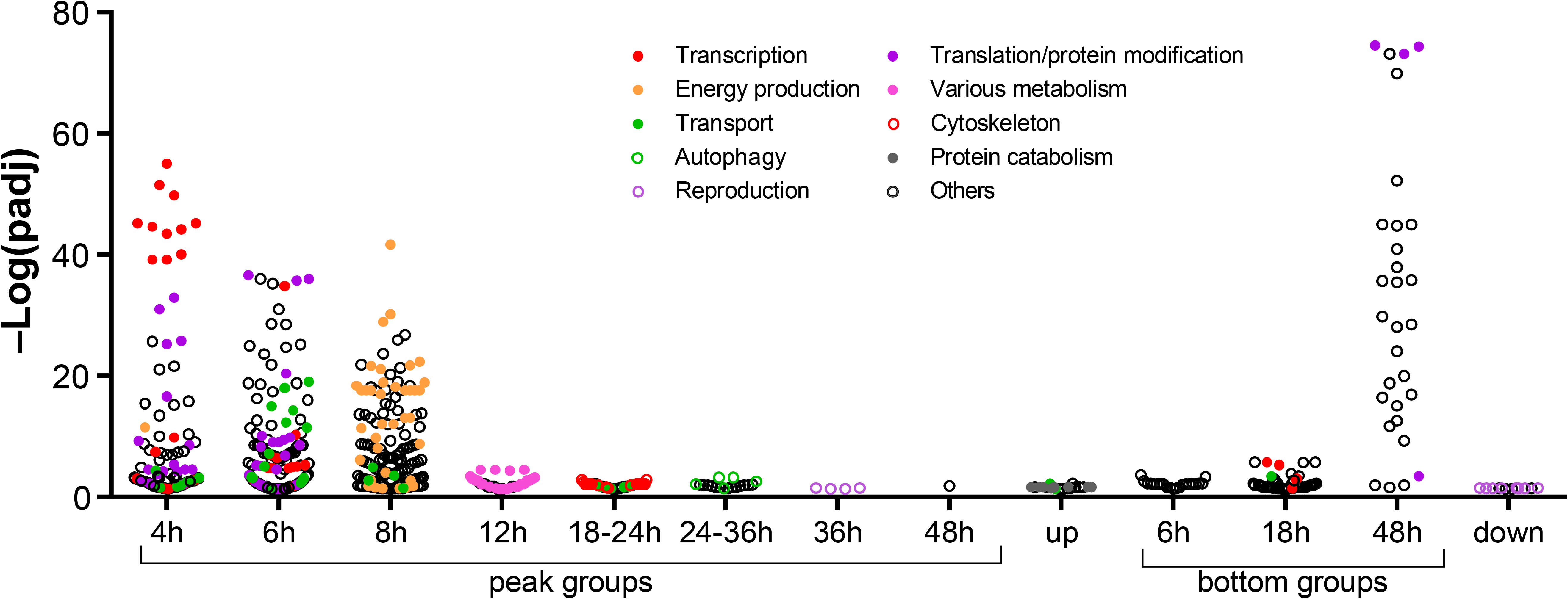
–log plot of Benjamini-Hochberg-adjusted p values (p < 0.05) for GO enrichment analysis (Biological Process) for each cluster. Colors are applied for the related terms discussed in the text, for example, red-filled circles are GO terms related to transcription.

The general trend of enriched terms for the peak groups showed the transitioning roles of the midgut as the center for bloodmeal digestion (Fig 4, Supplemental Table S2). The 4h peak group was enriched with GO terms for RNA metabolism, transcription, and translation, reflecting activation of transcription and protein synthesis. The largest change between time points shown in the PCA (1-4 hPBM) (Fig 2) may be due to this activation of transcription and translation, which suggests the female mosquito undergoes a drastic shift in their physiology to a digestion/reproduction state. The 6h peak group was found to be enriched with GO terms for translation (and protein synthesis), suggesting the system further shifted to protein production to deal with the bloodmeal. The 8h peak group was enriched with GO terms related to energy production, especially oxidative metabolism, suggesting a response to the high demand for energy needed to complete digestion. The 12h peak group was enriched with GO terms for a variety of metabolic processes such as lipid, carbohydrate and amino acid metabolism, oxidoreductase activity, and energy production, which are a potential reflection of the most active time for the bloodmeal digestion. The 18-24h peak group was enriched with GO terms for the regulation of cytoskeleton (mainly actin) and transport, implying that the midgut is arranging the cellular components to allocate the nutrients from the bloodmeal. The 24-36h peak group was enriched with GO terms associated with autophagy, suggesting recycling and regeneration of cellular components for recovery, may be at the transition of active digestion phase and recovery/regeneration phase. Although the 36h peak group was enriched with GO terms for oogenesis and reproduction, the genes represented by these terms were comprised of heme peroxidases (HPXs) and peptide hormone precursors, one of which is trypsin modulating oostatic factor (TMOF) (Borovsky 2003), which reduces the expression of trypsin in the midgut and in turn reduces oocyte maturation, suggesting a negative regulation of digestion in the midgut at the end of digestion, likely to shut down digestive processes. HPXs may act to suppress the midgut microbiome that had expanded upon bloodmeal, as HPXs mosquitoes appear to have immune functions (Kumar et al. 2010; Barletta et al. 2019). The 48h peak group was enriched with the genes with metal (mostly zinc ion) binding function and nucleic acid binding (“Molecular Function” terms, see Supplemental Table S2), which suggests modifying transcription by variety of DNA-binding proteins (transcription factors, enhancers, silencers, repressors, chromatin-modifying proteins, etc.). The “up” group was enriched with GO terms related to transport activity and ubiquitin-related functions, suggesting transport and protein turnover are important throughout the cycle as a hub for nutrients from the bloodmeal, implying the midgut as the center not only for digestion, but also for distribution. In addition, terms related to transport are present in other peak groups (Fig 3 and Supplemental Table S2), which also suggests that transport of nutrients are important in the midgut. The down-regulated gene groups (i.e. “bottom” groups) were enriched with GO terms similar to some peak groups, such as RNA metabolism (18h bottom, as in 4h peak), translation (48h bottom, as in 4h peak and 6h peak), reproduction (down, including peroxidases as in 36h peak). The 6h bottom group contained several terms with “regulation of” (Supplemental Table S2), which implies turning off the negative regulators of many biological processes. This may be due to role switch for the genes that have similar functions (genes for resting state vs active digestion state).

**Fig 4:**
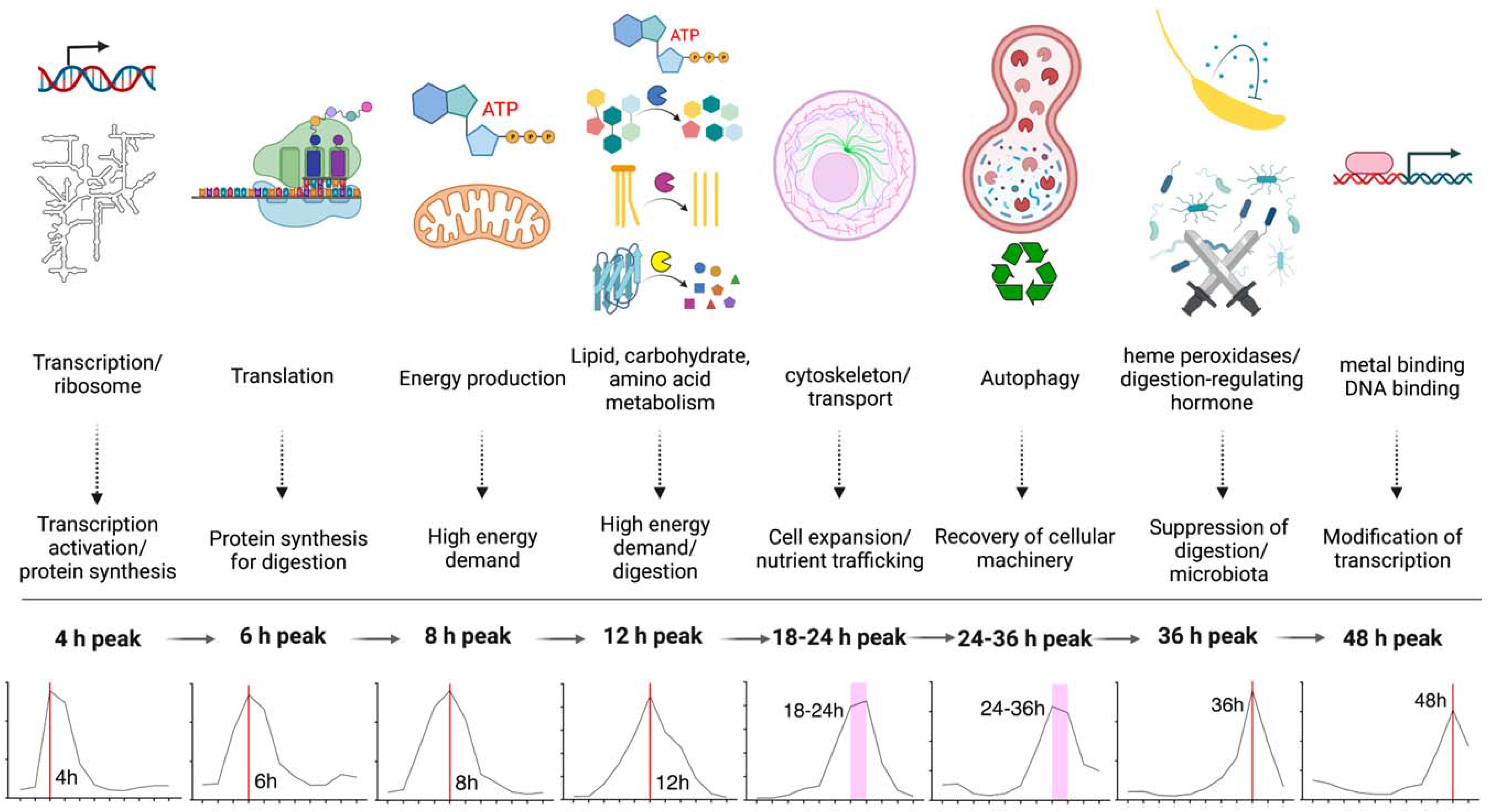
GO enrichment for the peak cluster groups (group 1-8). The descriptions below the graphics are directly related to the top GO terms. The description below the arrows are inferred roles. Created with BioRender.com

### Serine proteases

To highlight the utility of the temporal resolution provided by this dataset, we focused initially on a group of genes that have a key role in blood digestion and for which exists substantial prior work—serine proteases (SPs). With the midgut’s central role for bloodmeal digestion, SPs like trypsins and chymotrypsins have been described to play important roles in bloodmeal digestion in the midgut (Isoe et al. 2009). Indeed, throughout the time course, 8 out of the top 10 most abundantly expressed genes were SPs (Table 1) underscoring their importance in digestion.

**Table 1:**
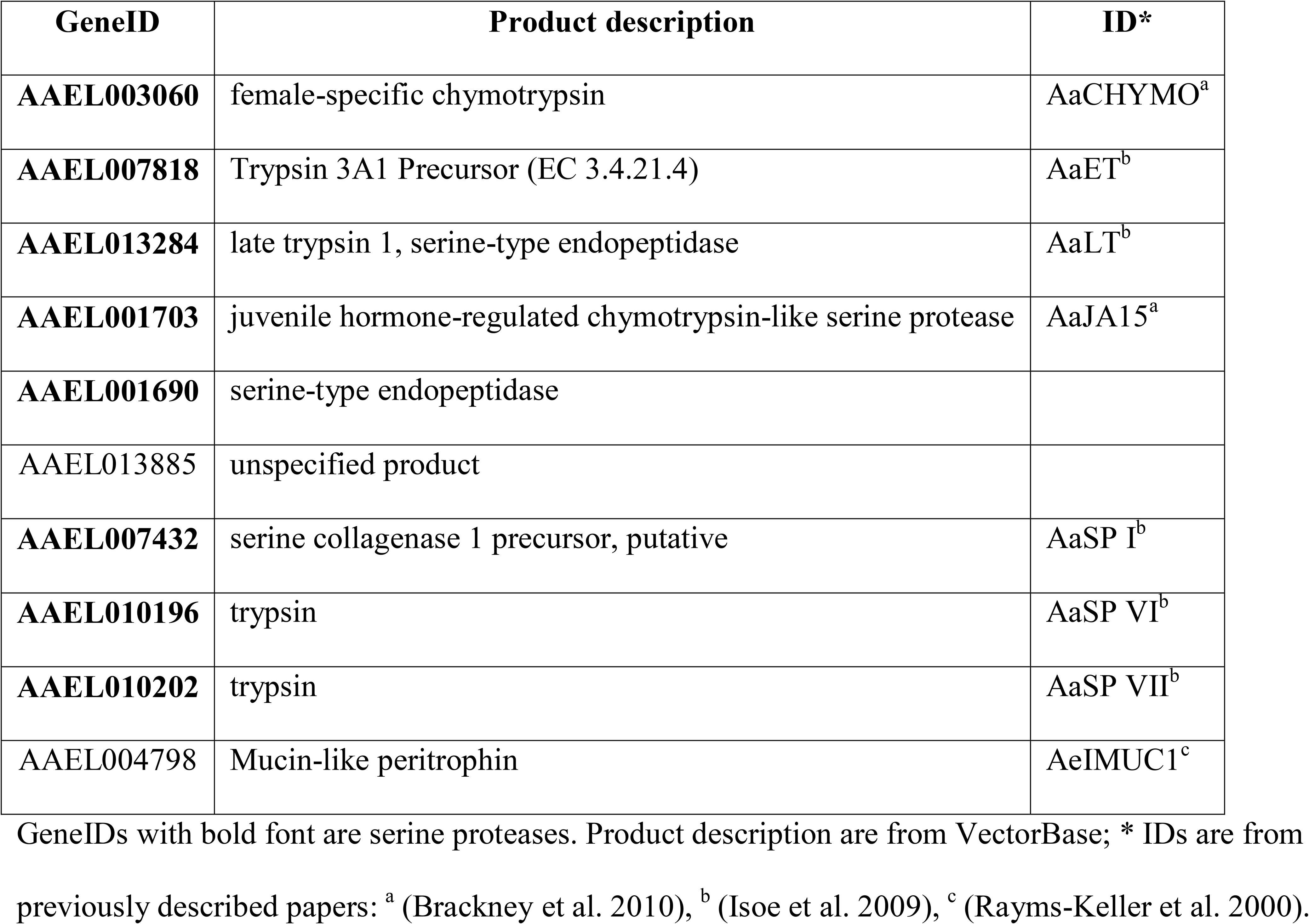
Top 10 most abundantly expressed genes in the midgut time-course transcriptome.

The top highly expressed gene, AAEL003060 (adult, female-specific chymotrypsin, AaCHYMO) showed very high transcript levels with a slight reduction trend over time (Fig 5), consistent with the observation by Jiang *et al*. that the expression of this protein is under post-transcriptional control (Jiang et al. 1997). The expression patterns of the early trypsin (AaET, AAEL007818) and the late trypsin (AaLT, AAEL013284) mirror each other, providing a more detailed view of the kinetics of the temporal compartmentalization of these major enzymes (Fig 5). While AaET transcripts were highest at 0 hPBM (pre-bloodfed) and <1 hPBM, Isoe *et al*. described that the protein was detected only at 3-6 hPBM (Isoe et al. 2009), suggesting post-transcriptional regulation occurs with this gene as well. AaLT transcript peaked at 24 hPBM, which corresponds the known protein expression pattern (Isoe et al. 2009). The top 10 list also includes a previously uncharacterized SP gene, AAEL001690. Expression of this gene is highly induced in response to a bloodmeal with a peak at 12-18 hPBM (Fig 5). Carboxypeptidase A (CPA), whose promoter has been the most widely used for midgut-specific transgene expression studies, is included for comparison.

**Fig5:**
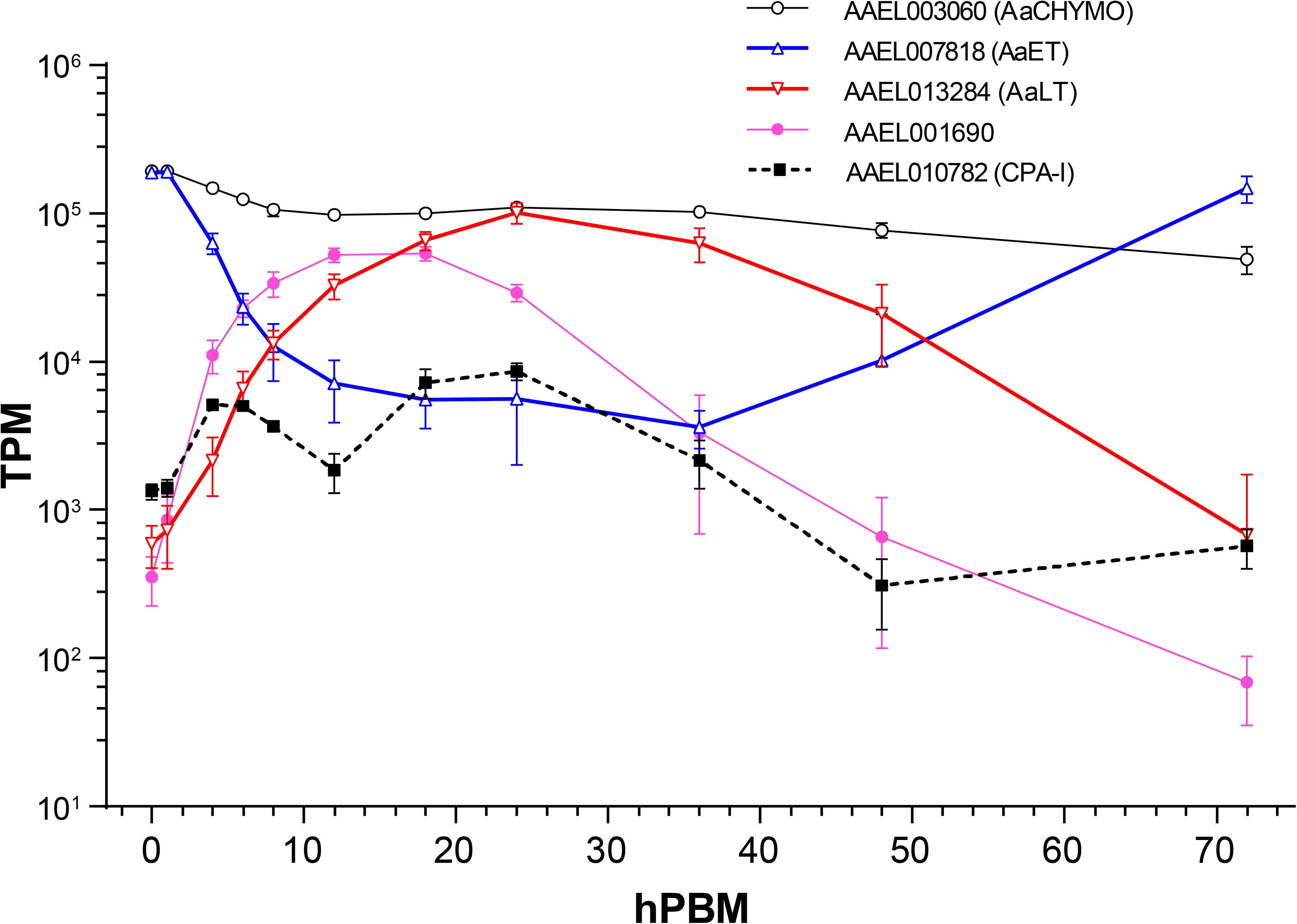
Expression of chymotrypsin (AaCHYMO, AAEL003060), early trypsin (AaET, AAEL007818), late trypsin (AaLT, AAEL013284), AAEL001690 and carboxypeptidase-I (AAEL010782, CPA-I, * not a serine protease). Error bars: SD

To get a comprehensive view of SPs expressed in the midgut, we obtained a list of all genes with the ontology term GO:0008236 (serine-type peptidase activity) in the *Ae. aegypti* genome at VectorBase. Out of 372 genes, we detected the expression of 346 genes in the midgut time-course transcriptome with non-zero TPM values (Supplemental Table S3), representing 93% of the serine proteases. However, since many of these genes showed very low expression, we limited the list of genes to those with log_10_(TPM+1) > 1.5 for at least one time point, which reduced the number to 56. A previous study by Brackney *et al*. described 12 midgut SPs (Brackney et al. 2010). Ten of the 12 genes were found in the current genome annotation (AaegL5.3) by nucleotide sequence similarity. The last two genes (AaSP II, GQ398044 and AaSP V, GQ398047) appear to have been omitted due to high nucleotide sequence similarity to AaSP III and AaSP IV, respectively, which implies that these pairs are allelic forms of the same genes or the genes remain to be found in the genome. Analysis of the expression patterns shows the presence of additional serine proteases that have expression patterns similar to those previously described (Fig 6). For example, AaLT (AAEL013284) exhibits a peak around 18-36 hPBM and the lowest at the 0-1 hPBM (as well as 72 hPBM). There are 11 other genes that show similar expression patterns (peak at 18-36 hPBM with lowest at 0 or 72 hPBM, orange shading in Fig 6) including some previously studied genes (AaSP VI, VII and Aa5G1) and 8 additional genes (also AAEL001690 discussed above) (Fig 6, orange shading), which may cooperatively participate in bloodmeal digestion. AaSP I (AAEL007432) shows elevated expression at earlier time points and remained high [log_10_(TPM+1) > 4] between 4 and 18 hPBM. There were 5 additional genes that exhibit similar expression patterns (Fig 6 purple shading), which may be important in initial digestion phase. Finally, there were also SP genes with an expression peak at later time points (36-48 hPBM, blue shading in Fig 6), suggesting potentially different roles in digestion. Thus, this study expands the existing knowledge about SPs in the midgut in context of the bloodmeal digestion physiology.

**Fig6:**
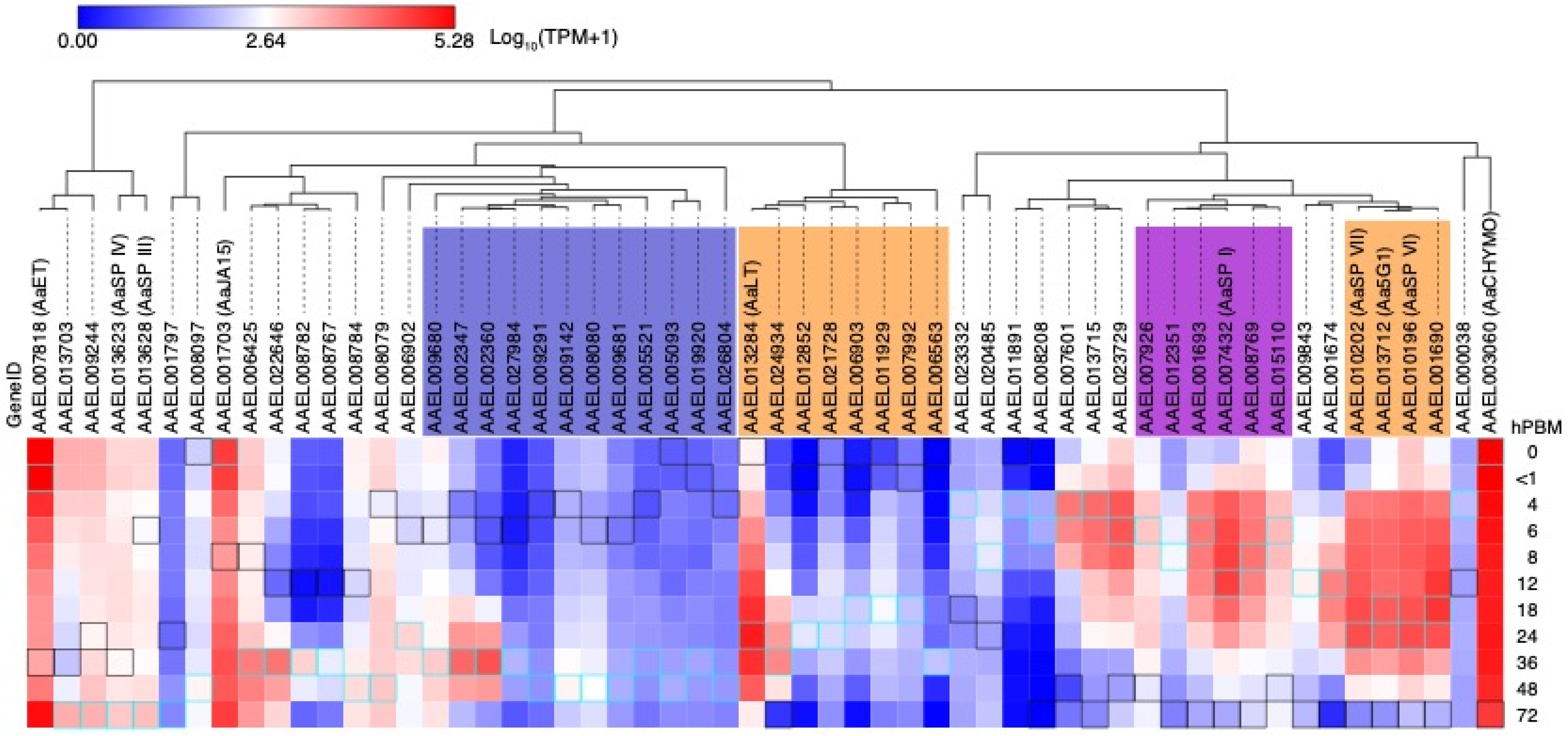
Expression patterns of serine proteases with 1.5 < log10 (TPM+1) at least one time point. Genes were hierarchically clustered by expression patterns. Maximum and minimum values across the time course are indicated by cyan and black squares, respectively. See text for the groups indicated by colored shadings.

A previous study indicated that targeting specific serine proteases by RNAi can slow digestion and reduce fecundity (Isoe et al. 2009) To confirm these results and investigate the role of these new SPs, we injected dsRNA targeting one or more SPs and evaluated the effects on reproductive output. We selected highly expressed SP genes in response to a bloodmeal, along with AaCHYMO, the most highly expressed gene, for two groups: the genes previously studied and showed reduced reproductive fitness by RNAi (AAEL013284 [AaLT], AAEL010196/AAEL013712 [AaSP VI/5G1; these genes were targeted/analyzed together as the sequences are nearly identical], and AAEL010202 [AaSP VII]) (Isoe et al. 2009), as well as AAEL001690, AAEL3060 [AaCHYMO], and AAEL007432 [AaSPI], genes that had not been previously characterized. Surprisingly, although we observed robust silencing of transcripts at 24 hPBM (Supplemental Fig S4) for all targeted genes, we did not observe any differences in fecundity (egg numbers) or fertility (hatch rates) (Supplemental Fig S5).

### Iron/heme related genes

Previously, we conducted screens for putative iron transporters (Tsujimoto et al. 2018; Tsujimoto et al. 2021), peritrophic matrix proteins (Whiten et al. 2018), and putative heme transporters (Eggleston and Adelman 2020) to better understand iron/heme handling from a bloodmeal in *Aedes aegypti*. Using a list of those genes along with ferritins, transferrins and recently determined heme exporters (Wang et al. 2022), we analyzed expression across our midgut transcriptome time course (Supplemental Table S4).

### Putative iron transporters (Fig 7 blue shading)

**Fig 7:**
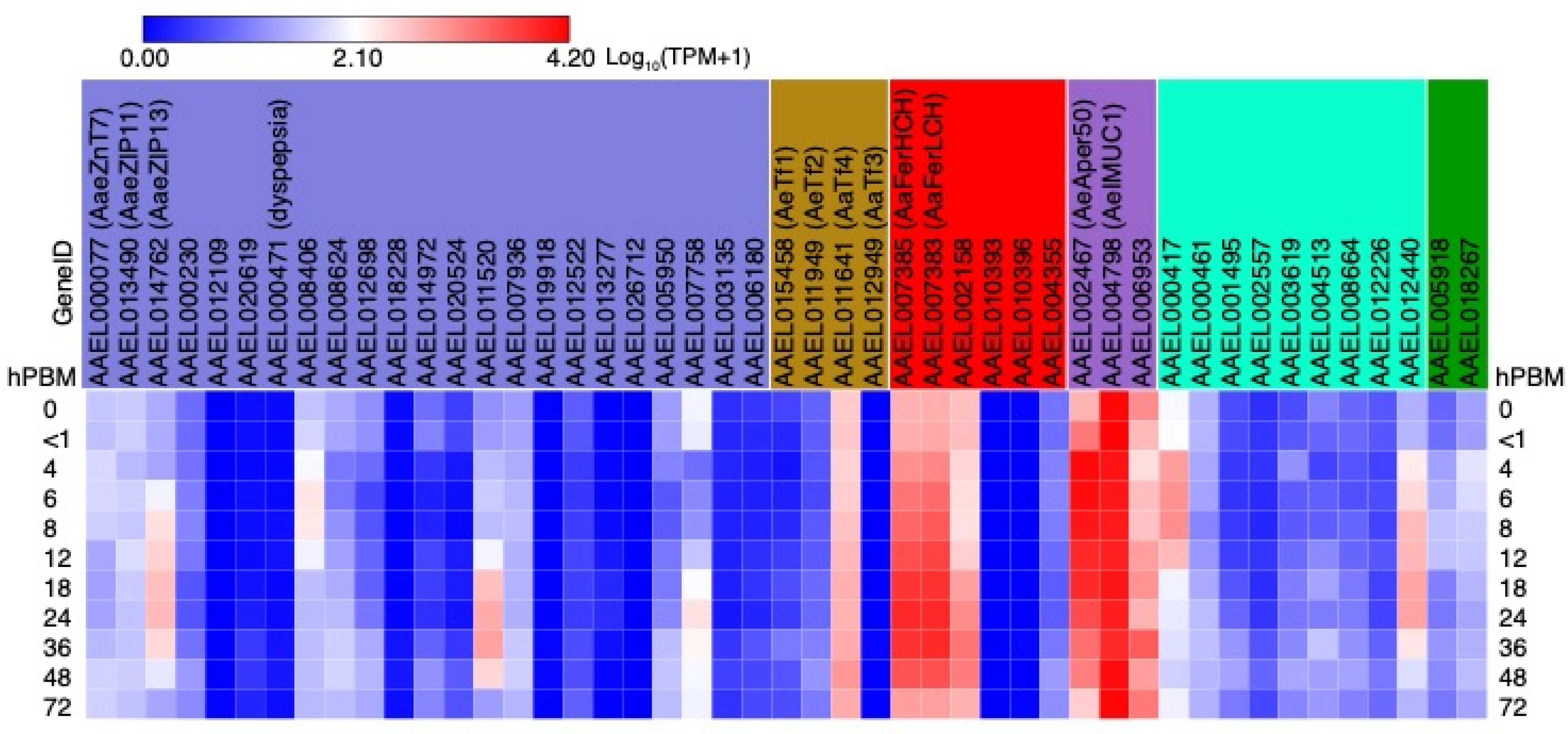
Heat map showing expression patterns of putative iron transporters (blue shading), transferrins (brown shading), ferritins (red shading), peritrophins (lavender shading), putative heme transporters (cyan shading), and verified heme exporters (green shading). The genes that showed no expression across all the time points were excluded. See also Supplemental Table S4.

Within ZnT and ZIP family putative iron transporters, AaeZIP13 showed elevated expression that peaked at 24hPBM (Fig 7), consistent with our previous observation (Tsujimoto et al. 2018). AaeZnT7 and AaeZIP11 showed relatively lower expression without drastic changes to transcript abundance across the time points, with just a moderate increase around 6 hPBM, 48-72 hPBM for AaeAZnT7, and 12 and 48 hPBM for AaeZIP11. Among genes identified in a screen for putative iron transporters, three (AAEL008406, AAEL011520, and AAEL007758) exhibited TPM higher than 100 (Fig 7, white to shades of red and Supplemental Table S4). Of these, AAEL008406 (annotated as a cationic amino acid transporter) displayed a peak of expression around 6-8 hPBM whereas AAEL011520 (SLC45/sugar transporter) and AAEL007758 (unknown function) showed peak expression around 24-36 hPBM (Fig 7 and Supplemental Table S4).

### Transferrins (Fig 7 brown shading)

Transferrins (Tfs) are known to be multi-functional iron-binding proteins (Zhou et al. 2009). Four Tf genes are present in the *Ae. aegypti* genome, among which two (AaTf1 and AaTf2) have been functionally studied. AaTf1 transcripts were found in fat bodies and adult female thorax, which were elevated by bloodmeal and bacterial challenge in the ovaries (Harizanova et al. 2005; Zhou et al. 2009). AaTf1 protein was also found in the hemolymph (Harizanova et al. 2005). AaTf2 transcripts are found in fat bodies and ovaries with suggested functions in development and wound response (Zhou et al. 2009). Here we show that unlike AaTf1-3, AaTf4 (AAEL011461) was highly expressed in the midgut ((Harizanova et al. 2005; Zhou et al. 2009) and Fig 7). Functional study on AaTf4 may uncover the role of AaTf4 in iron transport and control of gut microbiota upon bloodmeal.

### Ferritins (Fig 7 red shading)

Ferritins in general are known to be multimeric proteins that make a large sphere to cage thousands of Fe^3+^ (Pham and Winzerling 2010). Unlike mammalian ferritins, which serve as iron storage proteins, insect ferritins are considered to function as iron shuttle proteins and protectants from oxidative damages caused by Fe^2+^ (Pham and Winzerling 2010). The *Ae. aegypti* genome contains 7 ferritin-like genes, of which no transcript for AAEL000359 was detected in the midgut transcriptome (Supplemental Table S4). In *Ae. aegypti* ferritin light chain (FerLCH, AAEL007383) and heavy chain (FerHCH, AAEL007385) are well documented. Both proteins have been shown to be induced by a bloodmeal in the midgut (Dunkov et al. 2002). Transcripts of the two genes were induced by ferrous ammonium sulfate (Fe^2+^), hemin (heme-related compound), and H_2_O_2_ in whole bodies (Geiser et al. 2003). As expected, the most highly expressed ferritin genes in the midgut were AAEL007383 and AAEL007385 (Fig 7). These genes are closely located by head-to-head orientation, which was determined to contain multiple transcription start sites and documented FerLCH and FerHCH promoters (Geiser et al. 2003; Pham et al. 2003; Pham and Chavez 2005). The expression patterns and levels of these genes are nearly identical, which supports the idea that these genes are operated by a common promoter. Their increased expression after bloodmeal may be a response to elevated iron levels to mitigate the oxidative damage by Fe^2+^. The promoter for the AAEL007383 (AaFerLCH) was demonstrated to be specifically responsive to the intracellular iron levels in a cultured cell line, Aag2 (Tsujimoto et al. 2018). Another ferritin gene, AAEL002158, also exhibits high expression with induction at later stage (after 12 hPBM) than the other two, which implies a distinct role in bloodmeal digestion.

### Peritrophins (Fig 7 lavender shading)

All three previously described midgut peritrophins were found to be highly expressed across time course (Fig 7). AAEL004798 (AeIMUC1) exhibited very high expression throughout the time course; it is indeed one of the top 10 most expressed genes (Table 1), although its transcript expression was not induced by bloodfeeding. Since it has been demonstrated to bind a large amount of heme (Devenport et al. 2006), AeIMUC1 may be the major heme-sequestering peritrophin. AAEL006953 also showed varied expression levels across the time course. On the other hand, AAEL002467 (AeAper50) expression was highly induced by a bloodmeal, which agrees with previous report, although our study lacks the 2 hPBM time point, the potential true expression peak (Shao et al. 2005). The response was quite rapid; at <1 hPBM, TPM increased more than 3 times from 0 hPBM, reaching a peak (TPM: 13,025; 25× of 0 hPBM) at 4 hPBM and maintained at least half of the peak level until 18 hPBM (Supplemental Table S4). Intriguingly, protein expression of AeIMUC1 is post-transcriptionally regulated and exhibited peak at 4 hPBM (Devenport et al. 2006), the high transcript level prior to this time point may be prerequisite for the rapid production of high amount of protein product to protect the midgut. This also implies that at least AeIMUC1 and AeAper50 orchestrate their expression for a rapid formation of PM, which precedes the major digestive phase.

### Putative heme transporters (Fig 7 cyan shading)

Heme is a component of the major blood protein, hemoglobin. Like iron, heme can cause cellular damage despite also serving as a vital nutrient and a signaling molecule in trace amounts (Eggleston and Adelman 2020). In a previous study based on cell culture transcriptomes, several transporters whose expression levels were responsive to heme were identified (Eggleston and Adelman 2020). Among these, two genes exhibited high (TPM greater than 100) expression (AAEL000417 and AAEL012440) (Fig 7 and Supplemental Table S4) in our midgut transcriptome time course. Gene AAEL000417 (a member of the monocarboxylate transporter family) showed a rapid increase of transcript with a peak at 6-8 hPBM, while AAEL012440 (annotated as sodium-bile acid cotransporter) showed gradual increase with a peak at 18-24 hPBM. The fact that these genes respond transcriptionally both to heme exposure and to a bloodmeal in the midgut suggests potential roles in blood digestion/detoxification that remain to be determined.

Recently, a heme exporter has been characterized from *Drosophila melanogaster* (Wang et al. 2022). The same study verified that homologs in *Ae. aegypti* also had similar activity. The transcript expression of these genes (AAEL005918 and AAEL018267) in the midgut were not markedly high and were only mildly induced by bloodfeeding, suggesting either that very little heme is exported to the hemolymph or that there are additional heme exporters that remain to be determined that compliment this function.

### Comparison with Hixson *et al*. data

To highlight similarities and differences between this study and the recent time-course transcriptome study of the gut by Hixson et al. (Hixson et al. 2022), we compared peptidase genes shown in their Fig4F and found that our results are comparable (Supplemental Fig S6). In fact, despite the conclusion based on the PCA by Hixson et al., this result along with theirs support 72 h as the total length of the digestive cycle. We next compared expression patterns of 8 major serine proteases individually (Supplemental Fig S7). Expression patterns of these genes, in general, resemble between the two studies, just different in temporal resolution. Further we compared digestive enzymes and transporter genes expressed in the gut from Supplementary file 4 by their groupings: lipid digestion, lipid transporters, amylases, maltases & glucosidases, sugar transporters, peptidases, and amino acid transporters, representing by heatmaps (Supplemental Fig S8). This showed very comparable expression patterns across multiple groups of genes between the two studies. Thus, our study provides deep temporal resolution that can be overlaid on the spatial resolution provided by Hixson et al (2022).

## Discussion

This study utilized Illumina sequencing technology to present the midgut transcriptome from pre- bloodfeeding state to the completion of bloodmeal digestion at 72 h in 11 time points in the most important mosquito vector of arboviruses worldwide, the first study to cover a complete digestion cycle in *Ae. aegypti* with such fine temporal resolution. We obtained a total of over 1 billion paired-end fragments, which provided a depth of more than 26 million fragments per sample. The PCA showed a huge transcriptomic change between <1 and 4 hPBM, which suggests that additional time points between <1 and 4 hPBM are warranted in future studies, and may fill a useful knowledge gap in understanding midgut digestion physiology. The remaining differences in gene expression between 0 (start of the cycle) and 72 hPBM (end of the cycle; 1421 genes differentially expressed) may be due to the state of ovaries. Since we used gravid females, the midgut may require signals from non-gravid ovaries to revert to the pre-bloodmeal state, independent of the time elapsed. Thus, the PCA analysis along with our imaging results suggests that 72 h is indeed a complete (or close to complete) cycle of blood-digestion in the midgut.

Our PCA results differ from the recent results from Hixson *et al*. (Hixson et al. 2022), which suggested 48 h as a cycle. This discrepancy might be because this study used different strain of *Ae. aegypti* (“Liverpool” vs Thai strain), source of blood (citrated sheep blood vs live chicken), different tissue components (whole midgut vs crop, midgut, and hindgut combined), temperature (28 vs 29 °C), light cycle (14:10 vs unknown), and number of time points. Moreover, the PCA between the two studies might have used different components as principal factors. However, close comparison between the two studies revealed far more similarities than differences (Supplemental Fig S6-8). Thus, the time-course expression patterns between the two studies are reasonably similar, which suggests that the time-course study of both studies are comparable, although some experimental parameters were different. This also suggests that the *Ae. aegypti* Liverpool strain retains major digestion physiology similar to the “near” wild strain of *Ae. aegypti* (the Thai strain used in Hixson *et al*. study).

The clustering analysis and associated GO term enrichment analysis (Supplemental Fig S3, Fig4-4, Supplemental Table S4) showed that the midgut operates as the center for digestion (and transport) by regulating gene expression in an orchestrated manner to set the stage for the most important task for a female mosquito, reproduction. This analysis also suggested that the midgut functions as a distribution center for nutrients obtained from the bloodmeal.

The data we obtained may be utilized to analyze expression patterns of genes in the same functional category, such as the serine proteases (SPs) discussed in this manuscript, or other categories of interest such as secreted proteins, membrane-bound transporters, or immune pathways. Our analysis of SPs has revealed a number of previously undescribed SPs that may function synergistically or have different roles in bloodmeal digestion (Fig 6). The view of detailed temporal expression patterns helps categorize (or subcategorize) groups of genes, especially complex sets of genes. Even for a gene of interest in the context of the time course expression may provide us with a new insight, such as expression peaks that may be a component to consider when developing a control strategy (timing of application).

To investigate the utility of the data for functional studies, we conducted RNAi on selected SPs and assessed the effects on reproductive fitness. Unlike the similar study by Isoe et al., we did not observe reductions in fecundity or fertility in the gene-silenced mosquitoes despite the huge reduction of mRNA expression. This may be due to redundancy, as in each case there were additional SPs showing similar expression patterns (orange and lavender-shaded groups) that could compensate for the reduction of the targeted genes. Additionally, although RNAi substantially reduced transcript levels from their wild-type levels, even this residual remaining expression could have been sufficient to produce enough of the targeted protein, as suggested by their Ct values. Finally, as shown by our GO-term enrichment analysis of clustered co-expressed genes, the midgut substantially increases its metabolic activity, transcription and translation machineries upon bloodmeal acquisition (Fig 4, Supplemental Table S3), which might also compensate for a reduction of the SPs. Establishing the importance of various SPs may require the generation of stable, multiplexed CRISPR/Cas9-based gene edited strains to cope with the extreme redundancy built to protect the process of bloodmeal digestion.

In addition to bloodmeal digestion, an earlier study reported that the addition of a trypsin inhibitor in a DENV-containing bloodmeal reduced viral titer in the mosquitoes (Molina-Cruz et al. 2005), suggesting the importance of serine proteases in susceptibility to viruses. Bonizzoni *et al*., suggested that differences in susceptibility for dengue-virus (DENV) between different strains of *Ae. aegypti* may be related to a trypsin, particularly AAEL007818 (AaET). They detected a difference in reduction of AaET at 5 hPBM between DENV-susceptible and DENV-resistant strains (Bonizzoni et al. 2012b). However, our data suggest that SPs in the midgut microecosystem seem to be more complex and correlation of SPs to DENV infection and dissemination requires further investigation.

Despite the importance of the gut in both blood digestion and pathogen transmission, only two promoters have been empirically verified to date for the midgut-specific expression in *Ae. aegypti* : carboxypeptidase A-I (CPA-I) (AAEL010782) (Moreira et al. 2000) and AaET (Zhao et al. 2016). We confirmed that the CPA-I transcript expression has two peaks at 4-6 hPBM and 24 hPBM (Fig 5). Our observation parallels the study on transgene expressing luciferase under CPA-I promoter, particularly luciferase transcript expression (see Fig 4 of the reference), which exhibited strong signal at 8 hPBM and 20-24 hPBM with weaker signal at 12 hPBM (Moreira et al. 2000). Studies with particular interest in transgene expression in the midgut at 12 hPBM may consider avoiding the use of this promoter. AaET expression is robust pre-bloodmeal, but it reduced during the major digestion phase, confirming that the AaET promoter is not suitable for bloodmeal-inducible expression of transgenes. Promoters from genes like AaLT and AAEL001690 as well as the genes that showed similar expression patterns (orange groups in Fig 6) may serve for a more robust blood-meal inducible transgene expression in the midgut. Moreover, due to the high transcript expression, investigation of cis-regulatory elements (promoters) for the SPs may lead to future usage for midgut-specific transgene expression, although organ/stage-specific expression may need to be verified.

A recent study on transcriptome of all three tagmata (head, thorax, and abdomen) with subsectioned gut system and ovaries uncovered previously understudied spatial resolution of gene expression in the female *Ae. aegypti* (Hixson et al. 2022). This study complements their study by providing the finest temporal resolution known to date for bloodmeal digestion cycle of *Ae. aegypti*, which may serve as a platform for further studies. Moreover, similar studies based on the transcriptomics of other organs with fine time component like this should greatly enhance our understanding of physiology and genetics of bloodfeeding mosquitoes and lead to new research avenues.

## Materials and Methods

### Mosquitos

*Ae. aegypti* Liverpool strain was maintained in an insectary at Texas A&M Entomology in environmental chambers kept at 27 °C, 70% RH and 14:10 (L:D) cycles. Immature stages were reared with ground TetraMin (Tetra) and adults were fed on defibrinated sheep blood (Colorado Serum Company).

### Blood feeding and midgut dissection

For all the experiments, female mosquitoes 3-4 days post eclosion were used (the females were eclosed in the presence of males). They were maintained on 10% sucrose (granulated sugar from local grocery stores) in water and starved for ∼16 h prior to blood feeding. Mosquitoes were fed on citrated sheep blood (Hemostat Laboratories) via an artificial membrane feeder for ∼15 min, anesthetized on ice and only engorged ones were selected for midgut collection. Midguts were dissected from 0 (pre-bloodfed/starved, set aside when bloodfeeding was performed), <1 (no recovery from anesthesia), 4, 6, 8, 12, 18, 24, 36, 48 and 72 h post bloodmeal (PBM). The mosquitoes were discouraged from laying eggs on the sugar source (common with 10% sucrose) by increasing the concentration to 30% sucrose. To be consistent with the mosquito’s circadian rhythm, blood feeding was performed approximately 1 h after beginning of the light cycle. Blood meal (bolus) was removed from the midguts upon collection. A pool of 30 midguts were collected in a 1.5-mL microcentrifuge tube with TRIzol (Thermo Fisher) and stored at –80 °C for downstream processing. A total of 4 independent replicates were made for each time point.

### RNA extraction, library prep and sequencing

Whole RNA was extracted with TRIzol (Thermo Fisher) standard protocol with half volume format (0.5 mL TRIzol), and initial homogenization in a small volume of TRIzol (20 µL) using an RNase-free pestle (Contes). RNA pellets were resuspended with 16 µL of nuclease-free H_2_O and treated by DNase (2 µL of DNase buffer and 2 µL of TURBO DNase; Thermo Fisher) at 37 °C for 1 h. RNA was quantified by a Qubit RNA broad range kit (Thermo Fisher). Integrity of RNA was analyzed by a Fragment Analyzer (Agilent Technology).

Illumina libraries for RNAseq were prepared using NEBNext Ultra II RNA Library Prep kit for Illumina (E7770, NEB) with NEBNext Poly(A) mRNA Magnetic Isolation Module (E7490, NEB) and NEBNext Multiplex Oligos for Illumina (96 Unique index primer pairs) (E6440, NEB). We used 13 min fragmentation time and 9 cycles of PCR enrichment on 1000 ng of total RNA input determined by preliminary library preparation. All libraries were quantified by a Qubit with dsDNA BR kit (Thermo Fisher) and quality-checked by a Fragment Analyzer (Agilent Technology).

Pre-pooling quantification by qPCR, pooling and sequencing on a NovaSeq 6000 2 × 100 bp (paired-end) format, using an entire flowcell for all the libraries, were performed by the Genomics and Bioinformatics Service at Texas A&M Agrilife Research. The raw reads (fastq.gz) were deposited in National Center for Biotechnology Information (NCBI) Sequence Read Archive (SRA) with the BioProject ID: PRJNA639291.

### Bioinformatics analysis

The Illumina reads were mapped, and expression was quantified using HISAT2 (ver. 2.1.0) and StringTie (2.0) pipeline (Pertea et al. 2016) using the AaegL5 genome and AaegL5.2 geneset obtained from VectorBase (Giraldo-Calderón et al. 2015). StringTie was used with “-e” and “-A” options. Above processes were performed using the Texas A&M High Performance Research Computing system. A Count table (“gene_count_matrix.csv”) for downstream analysis (with DESeq2 and edgeR) was generated by a python script “prepDE.py” provided in the StringTie manual online (https://ccb.jhu.edu/software/stringtie/index.shtml?t=manual). A TPM (transcripts per million) table for each library was generated from count tables generated by the -A option of StringTie and the tables for all 44 libraries were combined into one table using a Python script. Pair-wise differential expression analyses and principal component analysis (PCA) were performed using DESeq2 package (ver. 1.34.0) (Love et al. 2014) on R (ver. 4.1.1). The combined TPM table was trimmed using a python script to remove the genes with 0 values across all the time points. The final TPM table is provided as Supplemental Table S5. and the gene IDs from the list were searched on VectorBase to obtain the information on the gene types (protein-coding, non-coding and pseudogenes).

Time-course analysis was performed following an example shown in edgeR package (on R ver. 4.1.1) (ver. 3.36.0) (Chen et al. 2016) user’s guide (June 2020 version), which uses smoothing spline with df=3. Through this, we obtained a list of genes showed significant change across the time course with false-discovery rate (FDR) < 0.05. Then using the FDR<0.05 gene list, we performed soft clustering analysis using an R package “Mfuzz” (ver. 2.54.0) (Futschik and Carlisle 2005) to obtain gene lists with similar expression patterns. With this analysis, we generated 20 clusters by expression patterns, which were consolidated into 13 groups. Gene Ontology (GO) enrichment analyses was performed for the Biological Process and Molecular Function categories using enrichment analysis option on VectorBase (https://vectorbase.org/vectorbase/app).

The codes used for HISAT2, StringTie, DESeq2 (including PHENO_DATA.csv table), edgeR, and Mfuzz are provided as Supplemental Material S2.

### Gut system and ovary imaging

Age-matched (to the mosquitoes used for sequencing) *Ae. aegypti* “Liverpool” females were bloodfed using a membrane feeder and defibrinated sheep blood. Midgut, Malpighian tubules, hindgut and ovaries were dissected at the same time points post bloodmeal. Unlike RNAseq samples, mosquitoes were bloodfed at different times of the day so that dissection could be performed in the working hours. The dissected organs were photographed using an AmScope MU300 digital camera (AmScope)-equipped stereo-microscope Leica M165FC. Scale bar was added by Fiji (ImageJ) (Rueden et al. 2017) using an image of a hemacytometer (Hausser scientific) grid with the same magnification as a scale reference.

### Serine protease gene family analysis

To obtain a list of serine proteases in the *Ae. aegypti* genome, we used the Gene Ontology term GO:0008236 (serine-type peptidase activity) on VectorBase. The expression data were retrieved from the master expression table (TPM) using a Python script. To filter SP with reasonable expression levels, the list was filtered to have log_10_(TPM+1) > 1.5 for at least one time point on Microsoft Excel (Microsoft). To visualize the expression patterns of these genes, Morpheus (https://software.broadinstitute.org/morpheus) was used using hierarchical clustering with default parameters (one minus Pearson correlation and average linkage). Modification of labels and dendrogram as well as high and low value highlighting were done using Illustrator (version 26, Adobe).

### RNAi analysis on serine protease genes

Primers for amplifying dsRNA templates were designed for serine proteases AAEL001690 and AAEL007432 (AaSP I) using Primer-blast (Ye et al. 2012) with PCR product size to be 350-600 bp and added 20 nt T7 promoter sequence at 5’ end. Resultant dsRNA sequences were searched against AaegL5.3 transcript set (from VectorBase) using standalone blast+ program (ver. 2.13.0) (Camacho et al. 2009) with “-word_size 19” option with no off-target was detected. Primer sequences for other genes (AAEL003060, AAEL013284, AAEL010202, AAEL010196/AAEL013712) were from (Brackney et al. 2008) and (Isoe et al. 2009) with some modifications (using 20 nt T7 sequence and 3’ deletion for annealing temperature adjustment). Phusion HiFi DNA polymerase (New England Biolabs) was used to amplify the template DNA, the products were purified using the Nucleospin Gel and PCR cleanup kit (Machery-Nagel), and quantified with NanoDrop One (Thermo Fisher). One _μ_g of the DNA was used as template for dsRNA synthesis using MEGAscript T7 Transcription kit (Thermo Fisher) at 37 °C overnight. The synthesized dsRNA was purified using MEGAclear Transcription Cleanup kit (Thermo Fisher), the resulting RNA pellets were resuspended with nuclease-free water, quantified by NanoDrop One and aliquoted for injection sessions. The dsRNA (∼1 _μ_g each target) was thoracically injected using Nanoject II (Dummond) into females within 48 h after eclosion. The females were kept with a cotton pad containing 10% sucrose solution. The cotton pads were removed ∼16 h prior to bloodfeeding. The starved females were bloodfed using artificial feeders and defibrinated sheep blood. After 15 min of bloodfeeding, the mosquitoes were anaesthetized on ice and only engorged ones were kept with a 30% sucrose cotton pad. Midguts were dissected at 24 hPBM (up to 7 individuals) for quantification of transcript (at least for a replicate) by qRT-PCR. At 72 hPBM, remainder were transferred to the EAgaL plates (Tsujimoto and Adelman 2021), which consist of 24-well tissue culture plates with agarose on the well bottom for fecundity and fertility evaluation. Briefly, the females were allowed to lay eggs for 24 h, and were removed from the EAgaL plates. Images of each well were captured by a compact digital camera (TG-6, Olympus), which were used for counting eggs using ImageJ (Rueden et al. 2017). Right after the well imaging, water was added to each well and monitored for hatching. Five days after oviposition, images of each well were captured for larval counting. GraphPad Prism (GraphPad) was used for statistical analysis (Kruskal-Wallis with multiple comparison test) and graphing.

Quantitative real-time PCR (qRT-PCR) primers were designed for AAEL001690, AAEL007432, and AAEL003060 using Mfold (Zuker 2003) for secondary structure prediction (at 60 °C) and Primer-blast with PCR product size to be 50-150 bp and 3’ self-complementarity “0”. Other primers were taken from previous studies (Brackney et al. 2008; Isoe et al. 2009). These primers were designed at least one primer to be outside of dsRNA site. qRT-PCR reactions were run on a Bio Rad CFX96 using SsoAdvanced Universal SYBR Green Supermix (Bio Rad). Amplification parameters were 95 °C for 30 s and 45 cycles of 95 °C for 15 s and 60 °C for 30 s followed by melt analysis between 65-95 °C. All reactions were performed in triplicate. Amplification efficiency of the primer pairs were empirically determined using serial dilution of midgut cDNA. % knockdown was calculated by the formula: % knockdown = (1 – [normalized expression of dsEGFP/normalized expression of dsTarget]) × 100.

All the primers with their information are shown in Supplemental Table S6.

### Comparison with Hixson *et al*., data

To show similarity and differences between this study and the study by Hixson *et al*. (Hixson et al. 2022), we retrieved Supplementary file 4 and used the TPM values from the file. We only used whole gut sugar fed, 4-6 (denoted 6), 24, and 48 hPBM data. TPM for the same sets of genes were extracted from our data (Table S1) using a custom python script. To generate a graph similar to Fig 3F in Hixson *et al*. the expression data were extracted from our master TPM table (Table S1), transformed the values to percent max across the time points for a better comparison using Excel. Line graphs were generated by plotting the percent max values using GraphPad Prism. Heatmaps were generated for Log_10_(TPM+1) values for both studies using Morpheus.

### Putative iron and heme transporters, and heme-binding proteins

The genes found to have relation to iron transport were obtained from (Tsujimoto et al. 2018) and from Table 1 in (Tsujimoto et al. 2021). The heme transporter candidates were from Table 6 in (Eggleston and Adelman 2020); heme exporters were from (Wang et al. 2022). The peritrophic matrix proteins with chitin binding domains were from (Whiten et al. 2018) (AAEL002495 is AAEL004798 in the current annotation AaegL5.3). Transferrins (Tf) and ferritins were retrieved by text search by “transferrin” and “ferritin” on VectorBase. The list of the genes are in Table S5. The expression values (TPM) for the genes were extracted from the master TPM table (Table S1) using a python script and log_10_(TPM+1)-transformed on Microsoft Excel. Morpheus (https://software.broadinstitute.org/morpheus) was used to visualize expression patterns after removal of any genes that had zeros for all the time points (AAEL019992 and AAEL000359). Additional labels and markings were made using Illustrator (Adobe).

## Data access

The raw reads (fastq.gz) were deposited in National Center for Biotechnology Information (NCBI) Sequence Read Archive (SRA) with the BioProject ID: PRJNA639291. All other data is available within the manuscript or as Supplemental Material.

## Acknowledgement

Fragment Analyzer quality analysis of RNA and the Illumina libraries and NovaSeq sequencing (including library quantification and pooling) were performed by the Texas A&M AgriLife Genomics and Bioinformatics Service. Portions of this research were conducted with the advanced computing resources provided by Texas A&M High Performance Research Computing. Authors also acknowledge Emma Jakes, Kristen Gilbert, and undergraduate research associates for helping with rearing, bloodfeeding and dissecting mosquitoes; Cara Muenzler for assistance for fecundity and fertility analysis (EAgaL plate assay).

## Author contributions

HT: conceived, designed, and conducted the project, analyzed and managed the data, wrote and revised the manuscript; ZNA: assisted with the experimental design and data analysis pipeline development and revised the manuscript.

