## Supplemental Material 1 for "An 11-point time course midgut transcriptome across 72 h after blood feeding provides detailed temporal resolution of transcript expression in the arbovirus vector, *Aedes aegypti*"

Supplementary Material 1:

Table of contents:


| | page |
|---|---|
| Supplemental Fig S1 | 2 |
| Supplemental Fig S2 | 3 |
| Supplemental Fig S3 | 4-5 |
| Supplemental Fig S4 | 6 |
| Supplemental Fig S5 | 7 |
| Supplemental Fig S6 | 8 |
| Supplemental Fig S7 | 9 |
| Supplemental Fig S8 | 10-15 |

Supplemental Fig S1

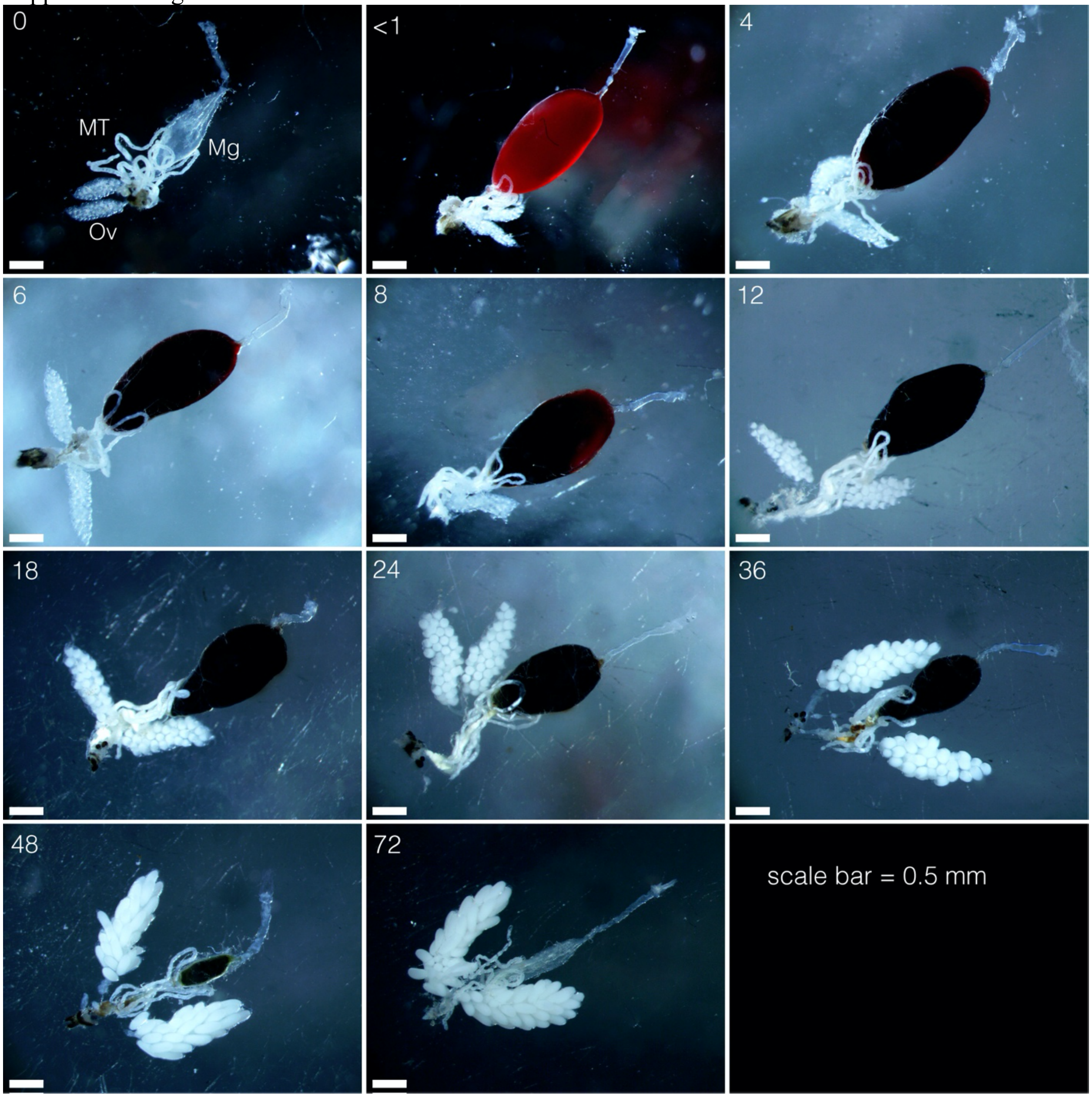


**Supplemental Fig S1**: Midgut with Malpighian tubules, hindgut, and ovaries dissected from age-matched females at time points sampled for the transcriptome study. Time points are indicated by numbers on the left top corner of each panel. Midgut (Mg), Malpighian tubules (MT), and ovaries (Ov) are labeled in the 0 hPBM panel. Hindgut locates between midgut and ovaries starting where MT arise mostly behind MT. Scale bar: 0.5 mm.

Supplemental Fig S2

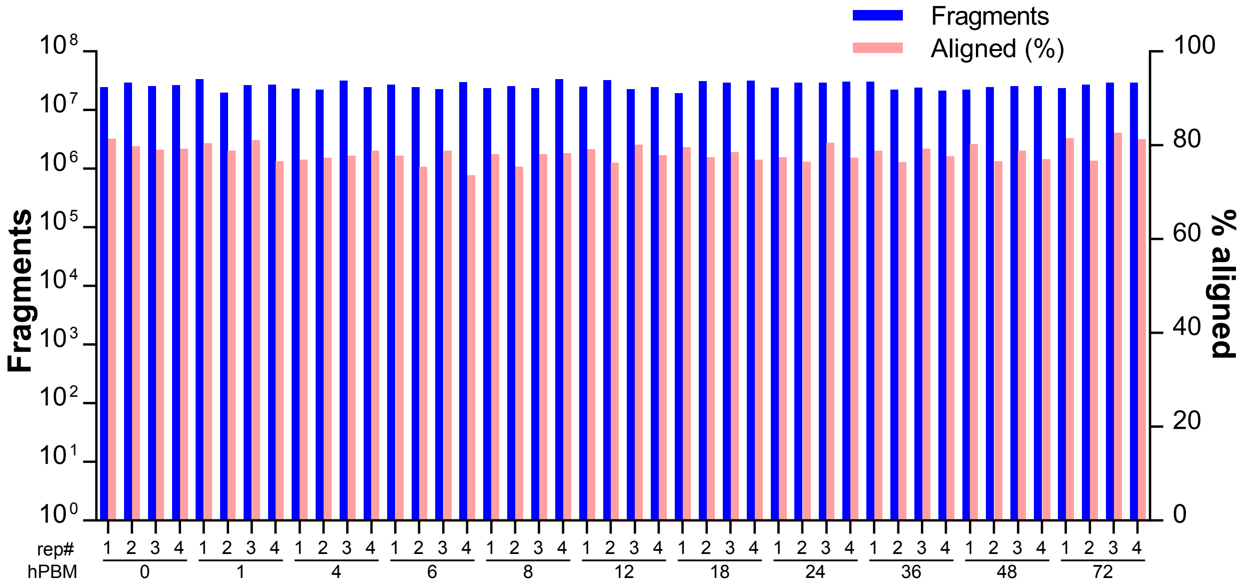


**Supplemental Fig S2**: Fragments obtained for the illumine libraries (blue bars, left vertical axis) and mapped fragments (pink bars, right vertical axis). Replicate numbers and time (hours) post bloodmeal (hPBM) are indicated on the horizontal axis.

Supplemental Fig S3

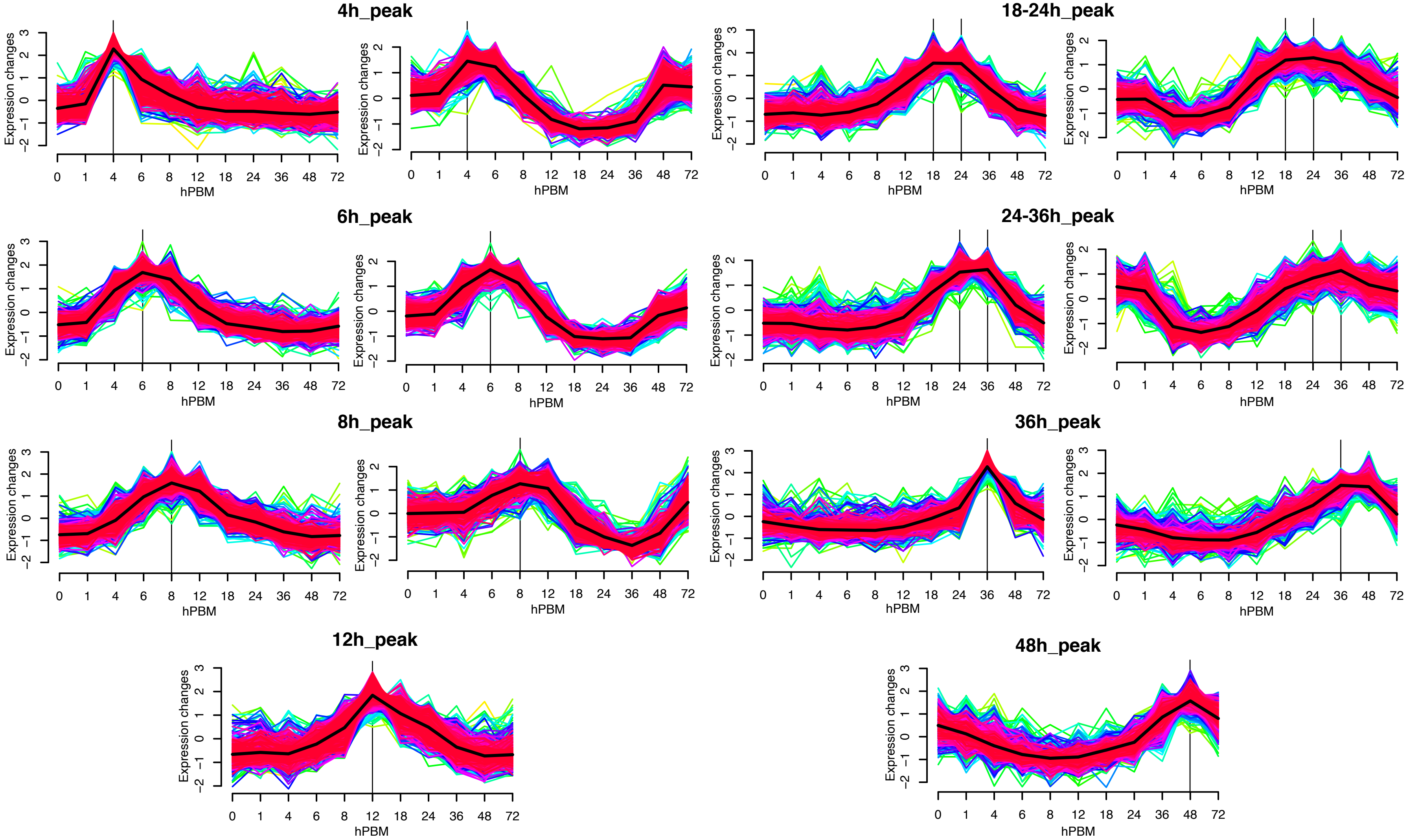

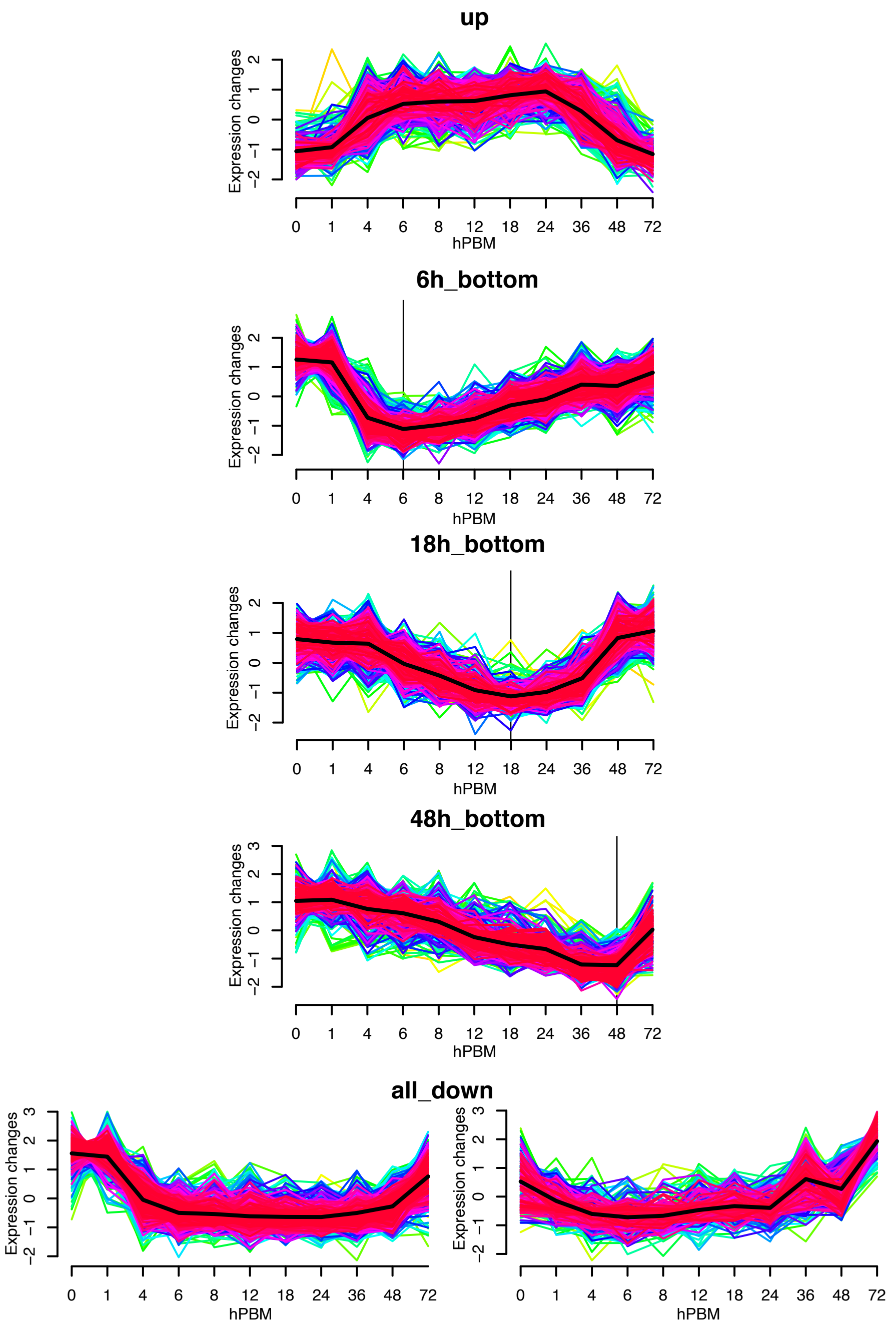


**Supplemental Fig S3**: Mfuzz soft clustering analysis. 20 clusters are grouped by similar expression patterns.

Supplemental Fig S4

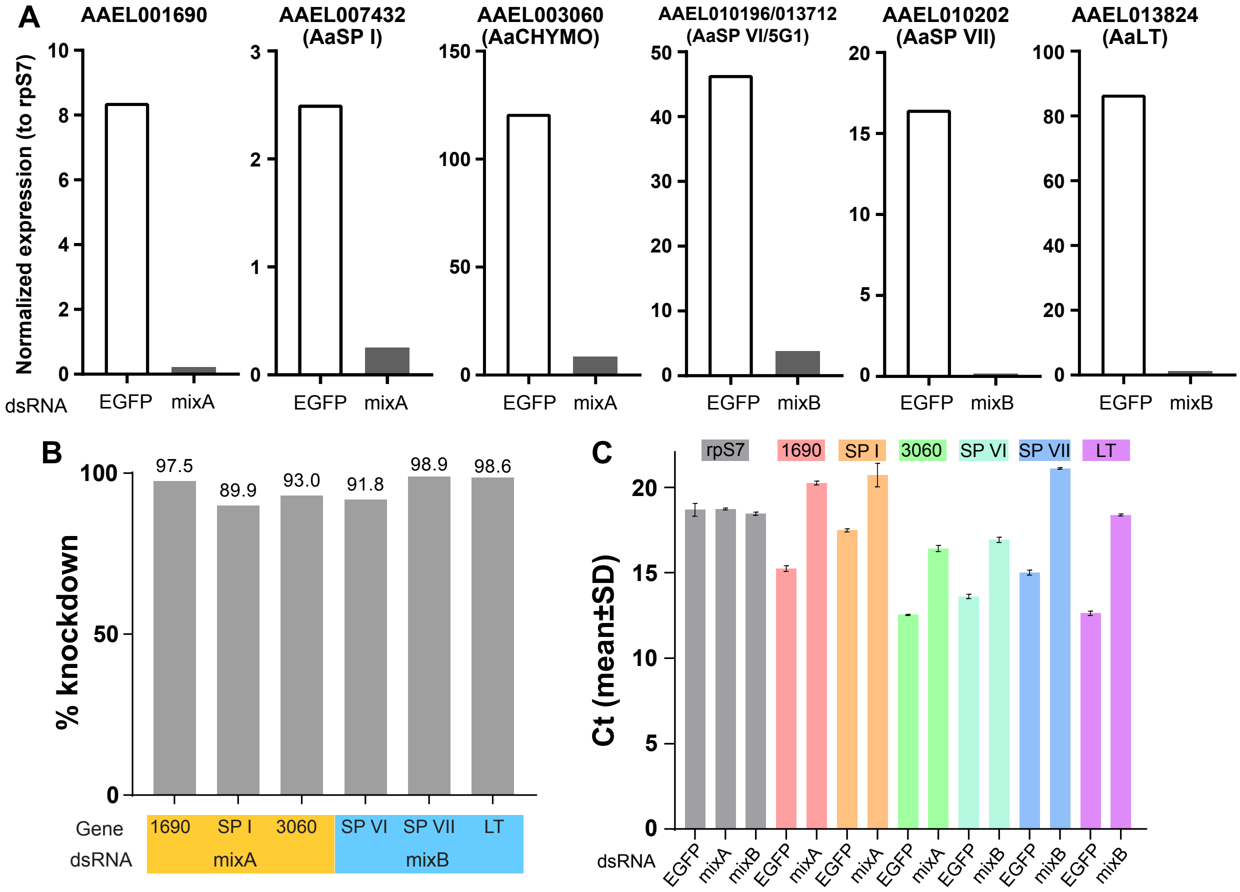


**Supplemental Fig S4**: Quantitative real-time PCR analysis to assess RNAi efficacy in the midgut 24 hPBM. **A** Normalized expression (to rpS7) between dsRNA against EGFP (control) and target gene-treated midgut. B Percent knockdown (in %) relative to EGFP control. C Raw Ct values (mean±SD for 3 technical replicates). mixA: mixture of dsRNA against AAEL001690, AAEL007432 (AaSP I), and AAEL003060 (AaCHYMO); mixB: mixture of dsRNA against AAEL010196/013712 (AaSP VI/5G1), AAEL010202 (AaSP VII), and AAEL013284 (AaLT).

Supplemental Fig S5

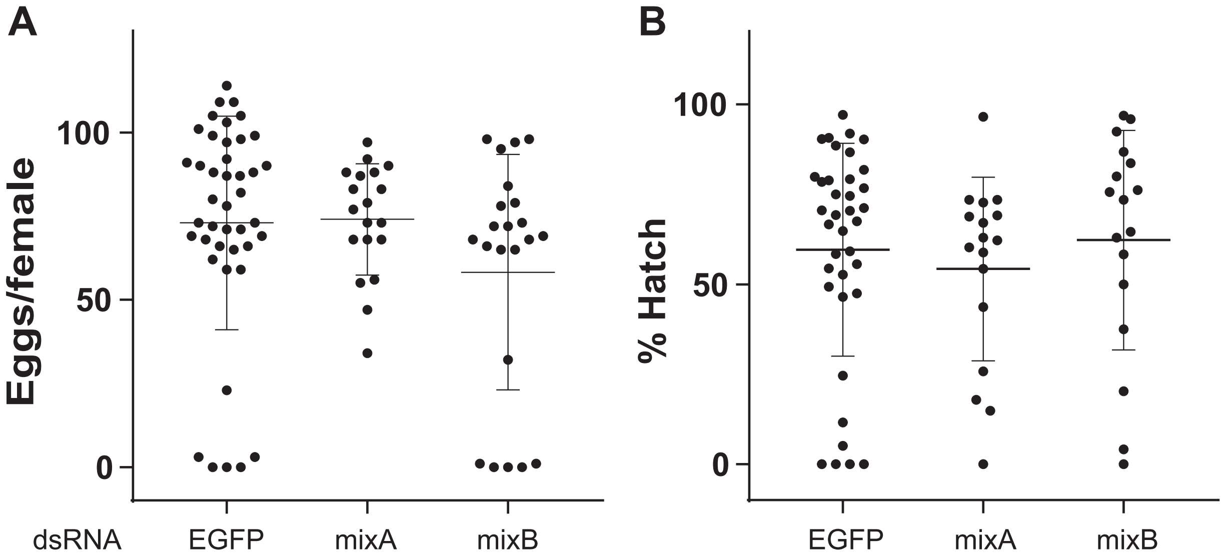


**Supplemental Fig S5**: **A** Fecundity (egg number) and **B** fertility (hatch rate) of dsRNA-injected females. mixA: mixture of dsRNA against AAEL001690, AAEL007432 (AaSP I), and AAEL003060 (AaCHYMO); mixB: mixture of dsRNA against AAEL010196/013712 (AaSP VI/5G1), AAEL010202 (AaSP VII), and AAEL013284 (AaLT).

Supplemental Fig S6

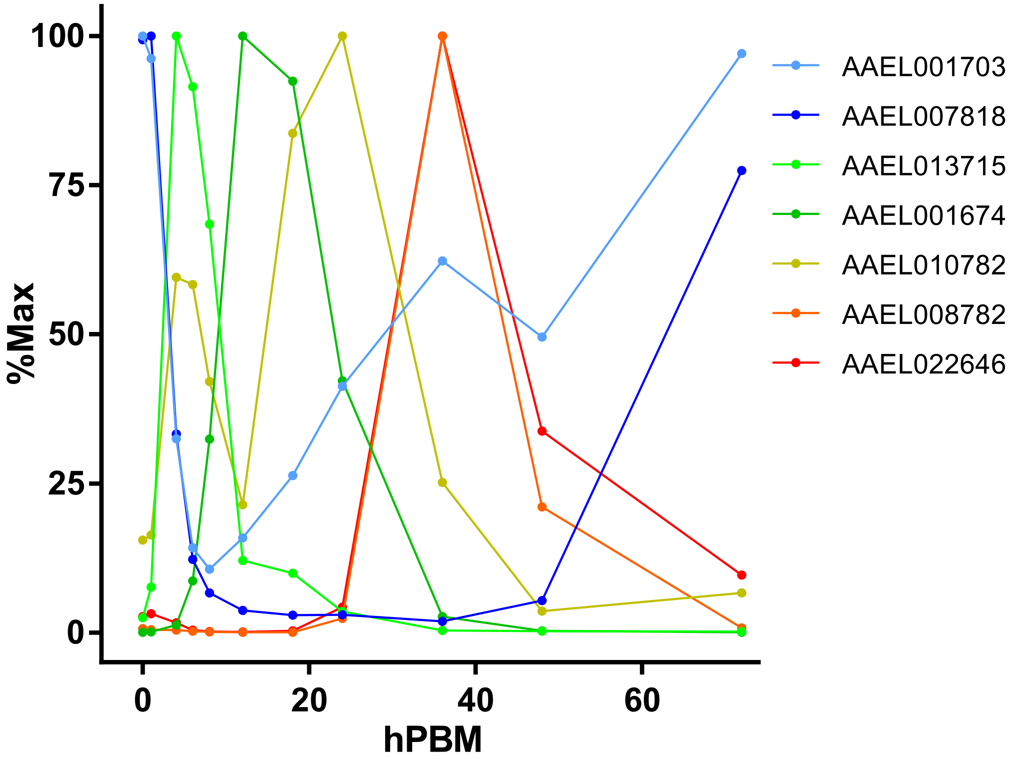


**Supplemental Fig S6**: % maximun (highest TPM for the gene as 100%) line plot for the peptidase genes presented in Fig 3F in Hixson *et al*., 2022.

Supplemental Fig S7

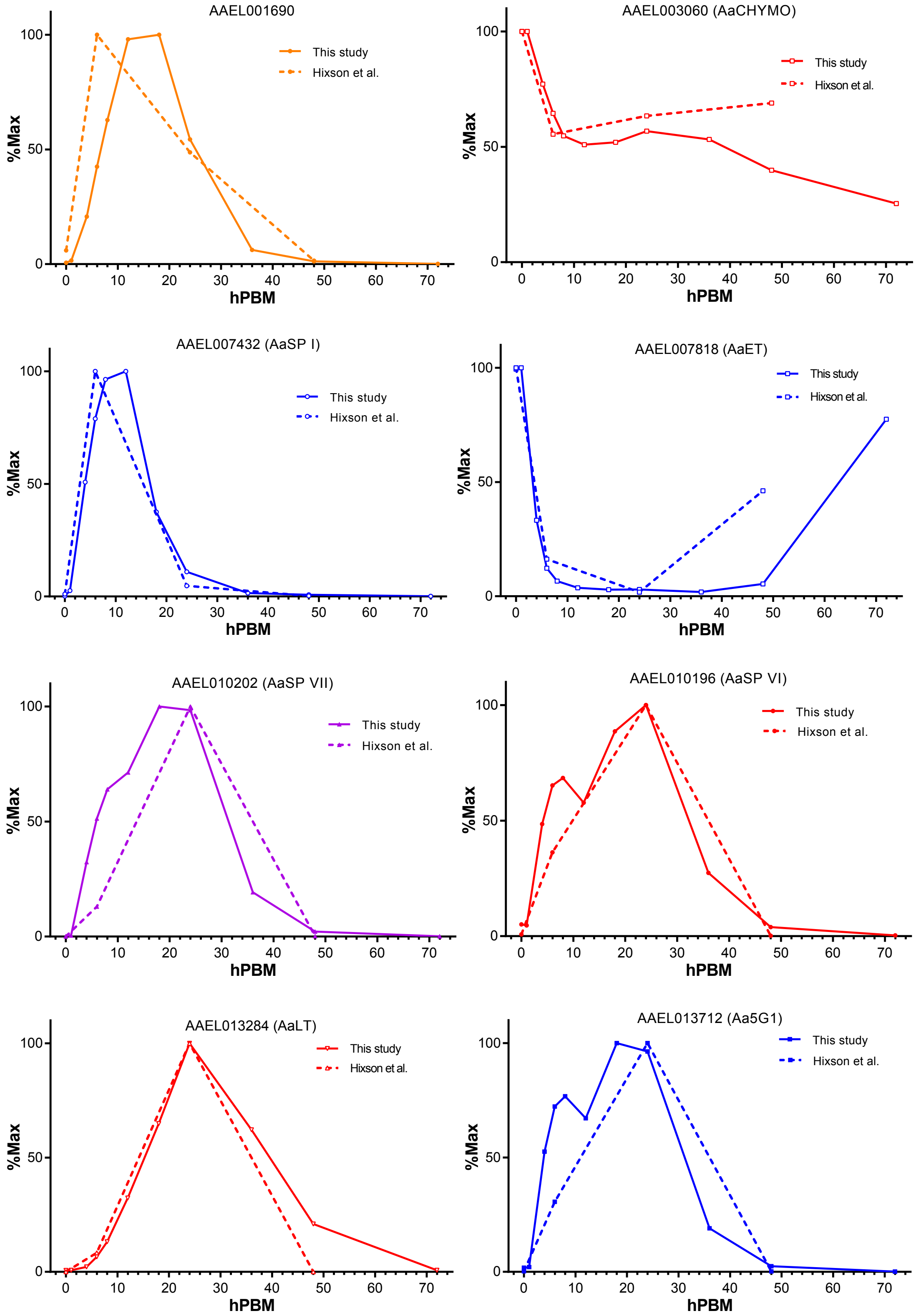


**Supplemental Fig S7**: % maximum (highest TPM for the gene as 100%) line plot for the peptidase genes to individually comparing expression of selected serine protease genes.

Supplemental Fig S8

#### (A) Lipid Digestion

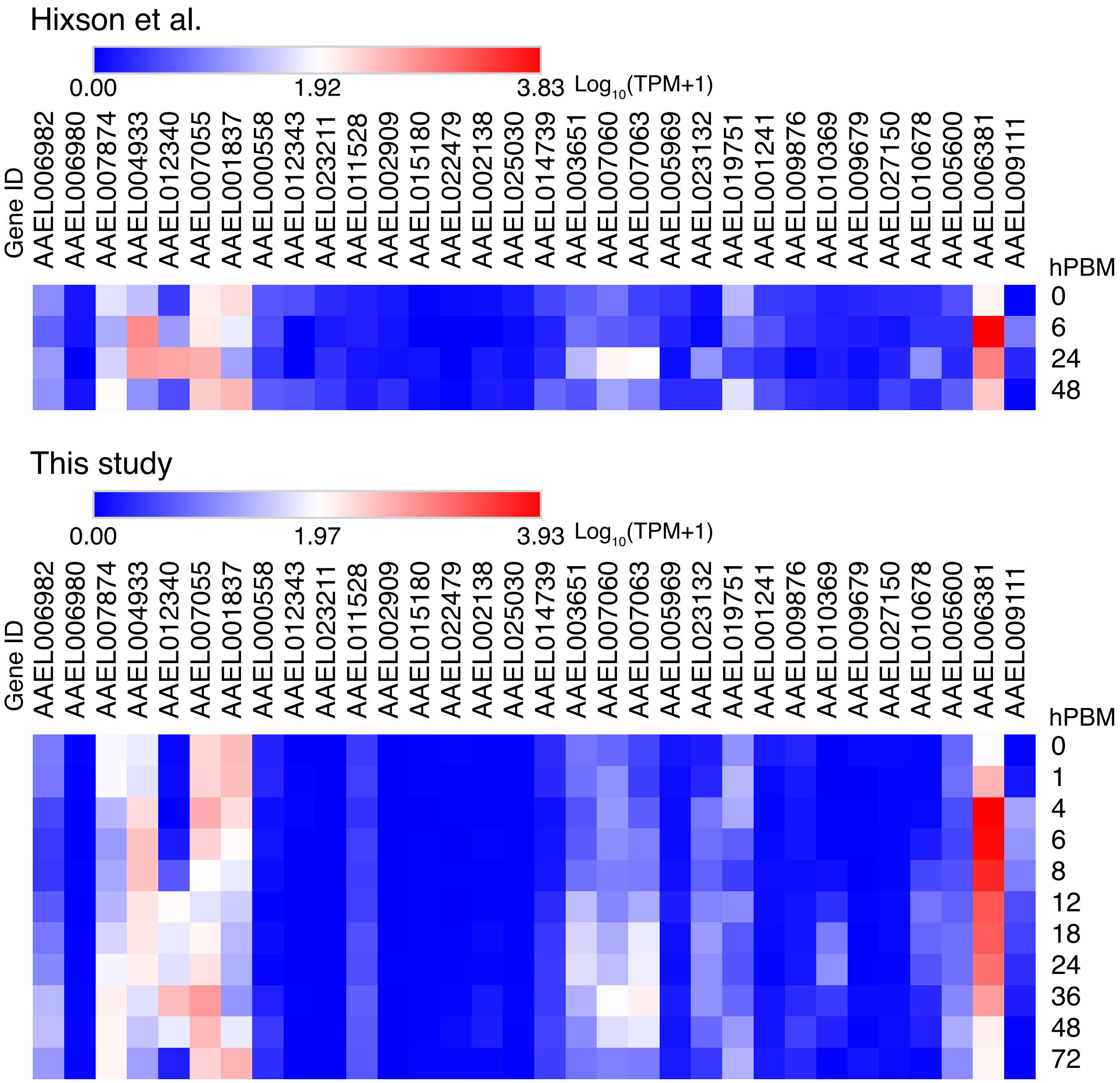

#### (B) Lipid Transporters

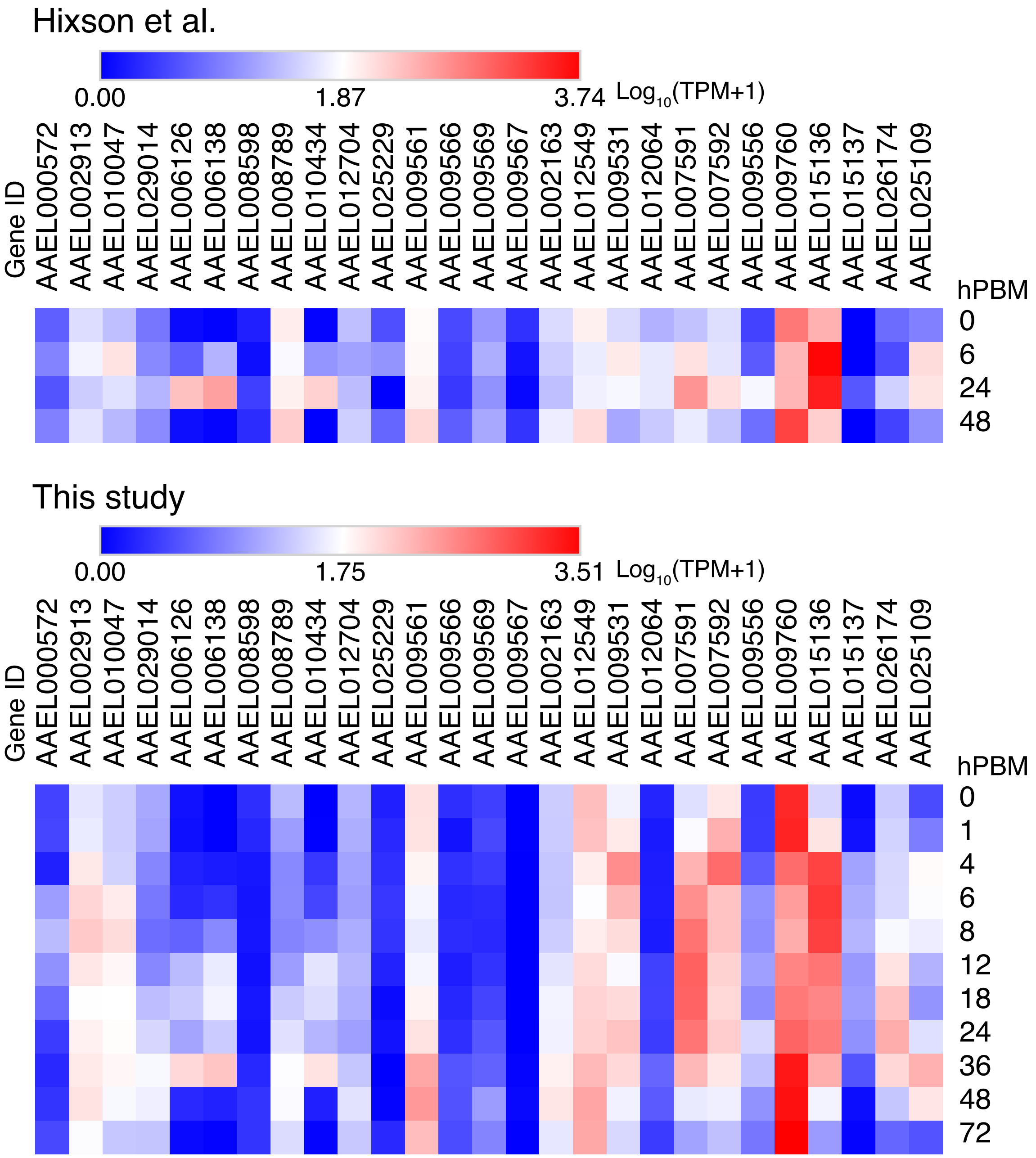

#### (C) Amylases, Maltases & Glucosideases

Hixson et al.

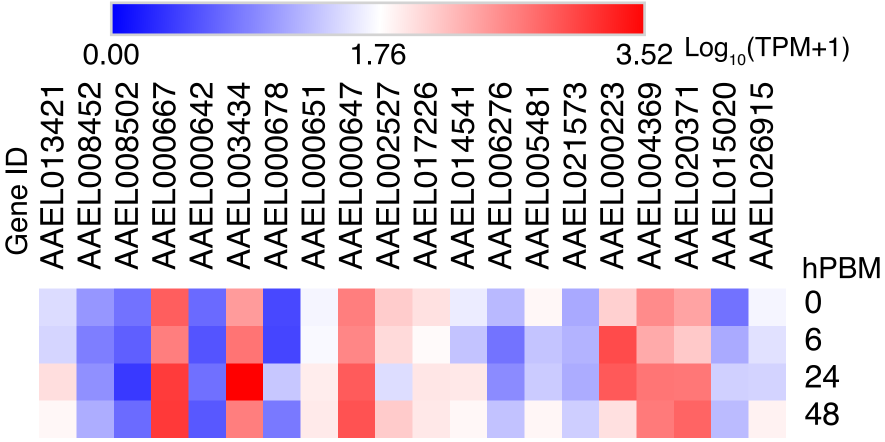


This study

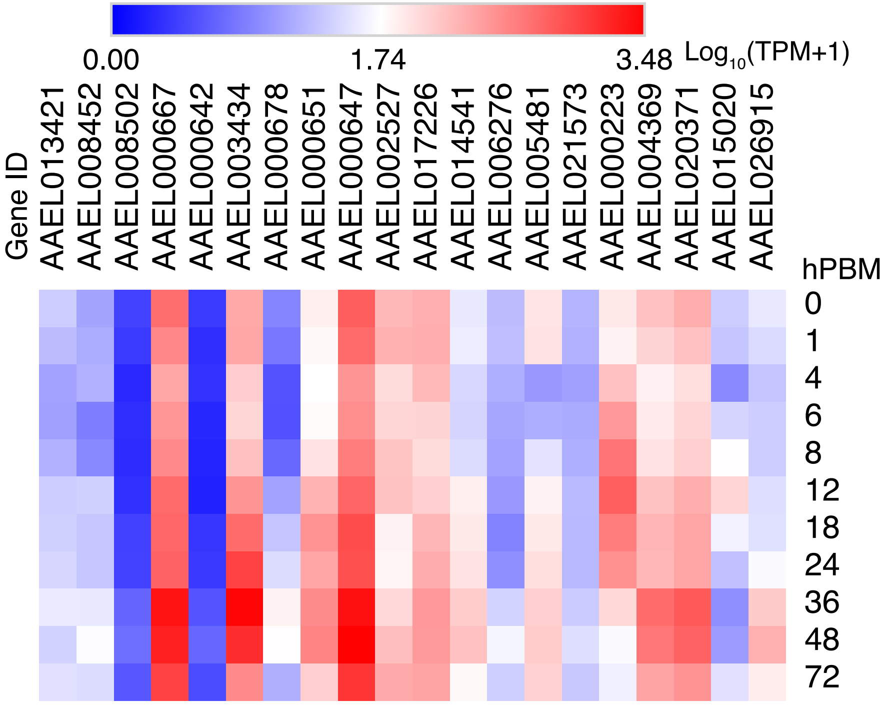

#### (D) Sugar Transporters

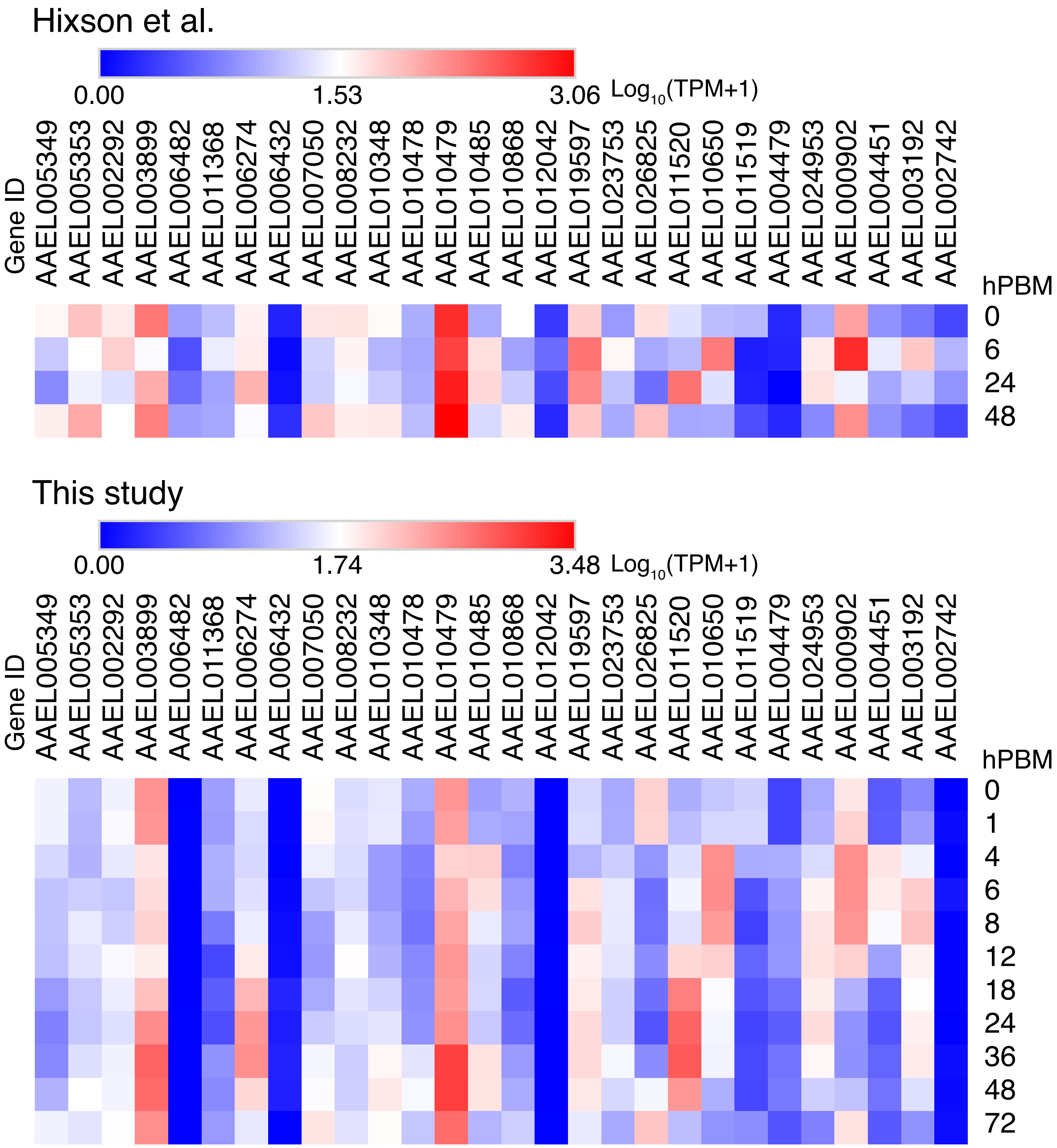

**(E) Peptidases**

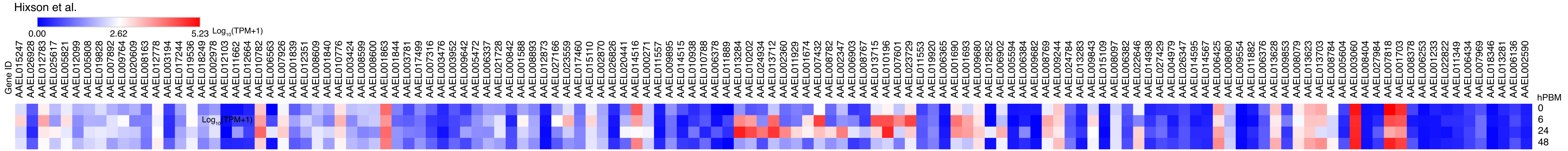


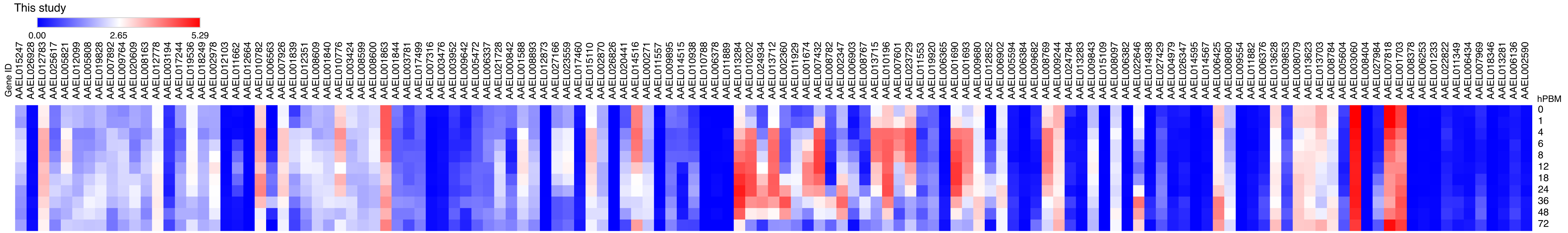

### (F) Amino Acid Transporters

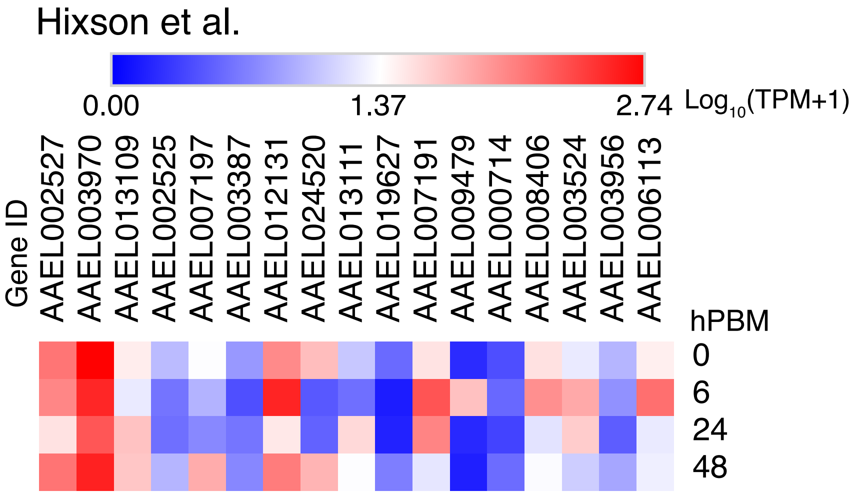


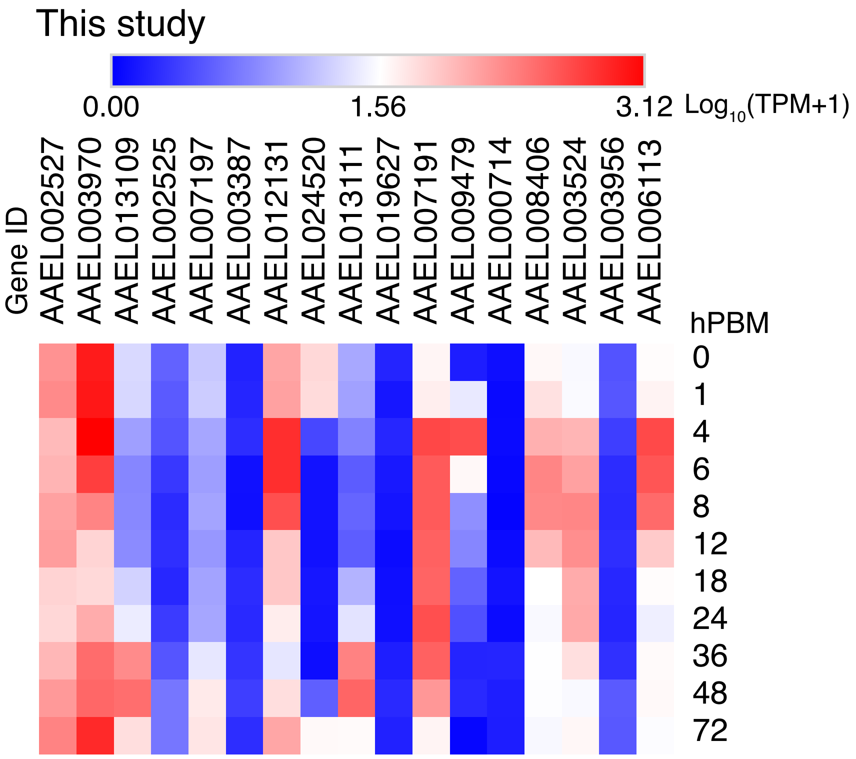


**Supplemental Fig S8**: Heatmap presentation of genes expressed in the gut in Hixson et al. (Supplementary file 4) by groups. A Lipid digestion, B lipid transporters, C amylases, maltases, glucosidases, D sugar transporters, E peptidases, F amino acid transporters.
